# Microtubule architecture connects AMOT stability to YAP/TAZ mechanotransduction and Hippo signaling

**DOI:** 10.1101/2025.08.08.669326

**Authors:** Giada Vanni, Anna Citron, Ambela Suli, Paolo Contessotto, Robin Caire, Alessandro Gandin, Giovanna Mantovan, Francesca Zanconato, Giovanna Brusatin, Michele Di Palma, Elisa Peirano, Lisa Sofia Pozzer, Carlo Jr. Albanese, Roberto A. Steiner, Michelangelo Cordenonsi, Tito Panciera, Stefano Piccolo

**Author notes:** Correspondence should be addressed to: Stefano Piccolo, Department of Molecular Medicine, University of Padua, IFOM, the FIRC Institute of Molecular Oncology viale Colombo 3, 35100 Padua, Italy. These authors jointly supervised this work: Tito Panciera, Stefano Piccolo.

## Abstract

Cellular mechanotransduction is a fundamental informational system by which cells read the structural features of their environment to control their own form and function. The YAP/TAZ transcriptional regulators are universal effectors of physical signals. Yet, the identity of proteins and subcellular structures serving as mechano-rheostats remains elusive. Here we demonstrate that perinuclear centrosome and microtubules’ architecture act functionally downstream of F-actin as cornerstones of cellular mechanotransduction. The mechanism revolves around the stability of AMOT proteins, that act as cytoplasmic inhibitory sinks for YAP/TAZ. Being degraded in mechano-activated cells and stabilized in mechanically-inhibited cells, AMOT serves as primary mechanical rheostat. In mechanically inhibited cells, microtubules form a cage-like network surrounding the nucleus, but, in mechanically activated cells, switch their architecture with formation of the centrosome from which microtubules sprout toward the cell periphery. In these conditions, AMOT proteins bound to the Dynein/Dynactin complex are subject to fast retrograde transport through the microtubular aster toward the pericentrosomal proteasome for quantitative and timely degradation. Restoring centrosomal condensation in mechanically inhibited cells by NLP1 overexpression is sufficient to restore mechanosignaling and YAP activation. AMOT proteins serves as universal integrator of distinct physical inputs from the ECM and the cytoskeleton, and their ablation renders cells invariably mechano-insensitive. Our findings also provide a unifying model that mechanistically merges mechanosignaling with the Hippo cascade. The current model by which YAP/TAZ are regulated by Hippo kinases is through direct YAP/TAZ phosphorylation. Our data instead show that, at least in the context of mechanotransduction, Hippo signaling inhibits YAP/TAZ largely indirectly, through LATS phosphorylation of AMOT averting it from its degradation route. We further show that Ras/RTK oncogenes hijack the AMOT degradation route to promote YAP/TAZ-mediated tumorigenesis. The findings imply the AMOT stabilization machinery as novel target for YAP/TAZ therapeutic modulation. In sum, our work reveals a previously unknown hierarchical coordination of distinct cytoskeletal and transport systems orchestrating mechanosignaling at the whole cell level.

## Main

Mechanical forces are increasingly recognized as pervasive factors in cell physiology, controlling multiple aspects of cell behavior, including differentiation, proliferation and tumorigenesis ^1–4^. The ability to sense mechanical forces must rely on the cell’s own construction principles: physical and architectural properties of the microenvironment, initially perceived at adhesive sites, need to be transmitted to the cytoskeleton and then other, still poorly understood, intracellular transducers. The discovery of the YAP/TAZ transcriptional regulators as pervasive transducers of physical signals into gene-expression programs opened the possibility to mechanistically dissect and functionally probe cellular mechanosignaling ^5, 6^. Yet, in spite of extensive efforts, it remains largely enigmatic how mechanically-induced changes in the cell’s cytoskeleton may ultimately impact on a dominant event of mechanotransduction: displacement of YAP/TAZ from so far poorly understood cytoplasmic anchors, allowing YAP/TAZ entry into the nucleus ^4, 7^.

In this view, a mechanical continuum of interconnected subcellular systems must be in place to allow the cell to entirely restructure itself in response to mechanical cues. Major emphasis in cell mechanics has been so far placed on F-actin microfilaments, whose structural organization and tensional state is directly connected to cellular adhesive systems ^8, 9^. Dissolution of F-actin filaments by drug treatments is known to prevent YAP/TAZ nuclear accumulation; conversely, boosting microfilament formation, as in cells lacking F-actin severing proteins, potently drives YAP/TAZ activation ^5, 10^. Yet, the direct link between YAP/TAZ control and F-actin still largely relies on inference. In fact, no biochemical mechanism involving bona-fide F-actin regulated protein has been directly connected to YAP/TAZ control^6^.

With this background in mind, we started this investigation by considering the contribution of cytoskeletal systems other than F-actin in connecting cell shape changes to mechanotransduction. Besides their well-established function in cell division, microtubules (MTs) are fundamental regulators of cell shape and motility, *on par* with actin-based processes ^11–14^. MT-associated proteins have been recently reported to tune Focal Adhesions at the cell periphery ^11^. Yet, how MTs switch their own architecture in response to distinct physical cues and their direct involvement in mechanosignaling is only rudimentarily understood. This represents a potentially large black box, as MTs are a self-organizing, pervasive and highly dynamic system, whose spatial organization (e.g., presence or absence of the centrosome or its location) is known to be intertwined with distinct cell states and functions ^15^. Just like F-actin, MTs connect to the nucleus, whose role in mechanotransduction has recently emerged ^16, 17^. Crucially, MTs’ architecture spatially organizes several subcellular structures by coordinating protein and organelle trafficking, in turn preserving cellular structure and function ^15^. By focusing on MTs, here we shed new light on the mechanisms of mechanosignaling; in so doing, we advanced on the view that the response to mechanical cues is coordinated at the whole cell level, involving a hierarchical coordination between distinct subcellular structures.

## Results

### Microtubule centrosomal organization as dominant determinant of mechanosignaling

We adopted MT-compatible immunofluorescence (IF) procedures, and MT-specific live imaging probes to investigate the structural organization of the microtubule lattice in cells either seeded on soft 0.7 kPa ECM or confined to small adhesive areas (hereafter mechano-OFF cells), and compared them to cells either stretched on a rigid 40 kPa ECM or on unconfined substrates (mechano-ON cells). Mechano-ON cells displayed a prominent perinuclear centrosome, labelled by γ-tubulin and pericentrin, established markers of the Microtubule Organizing Center (MTOC) (Fig. 1a-c, Extended Data Fig. 1a,b and Supplementary Video 1). As expected ^15^, MTs sprout in an astral arrangement from the MTOC, radiating toward the cell periphery, displaying a planar polarized alignment along the cell elongation axis (Fig. 1a- c, Extended Data Fig. 1a,b and Supplementary Video 1). Strikingly, however, MT organization dramatically changes in response to mechano-inhibitory inputs: mechano-OFF cells lose the MTOC, and their MTs become acentrosomal and display an inverted polarity, with minus-ends anchored to multiple peripheral γ-TURCs, mainly located beneath the apical cell cortex; all in all, this generates a MT network with an apico-basal “cage-like” structure around the nucleus (Fig. 1a-c, Extended Data Fig. 1a,b and Supplementary Video 1). These findings were obtained independently in different cell types, such as MCF10A, HEK293 and U2OS cells (Extended Data Fig. 1c,d), and show that comparable MT restructuring occurred in distinct experimental mechano-ON vs. -OFF conditions, that is, in cells plated on stiff vs. soft substrates, on large vs. small adhesive islands, in sparse vs. high density, or in cells experiencing mechanically dynamic hydrogels ^18^ in absence of cell detachment. In sum, changes in MT spatial organization are intrinsic to the cellular response to mechanical cues, with mechano-ON, but not mechano-OFF cells, displaying a perinuclear MTOC and centrosome.

**Fig. 1:**
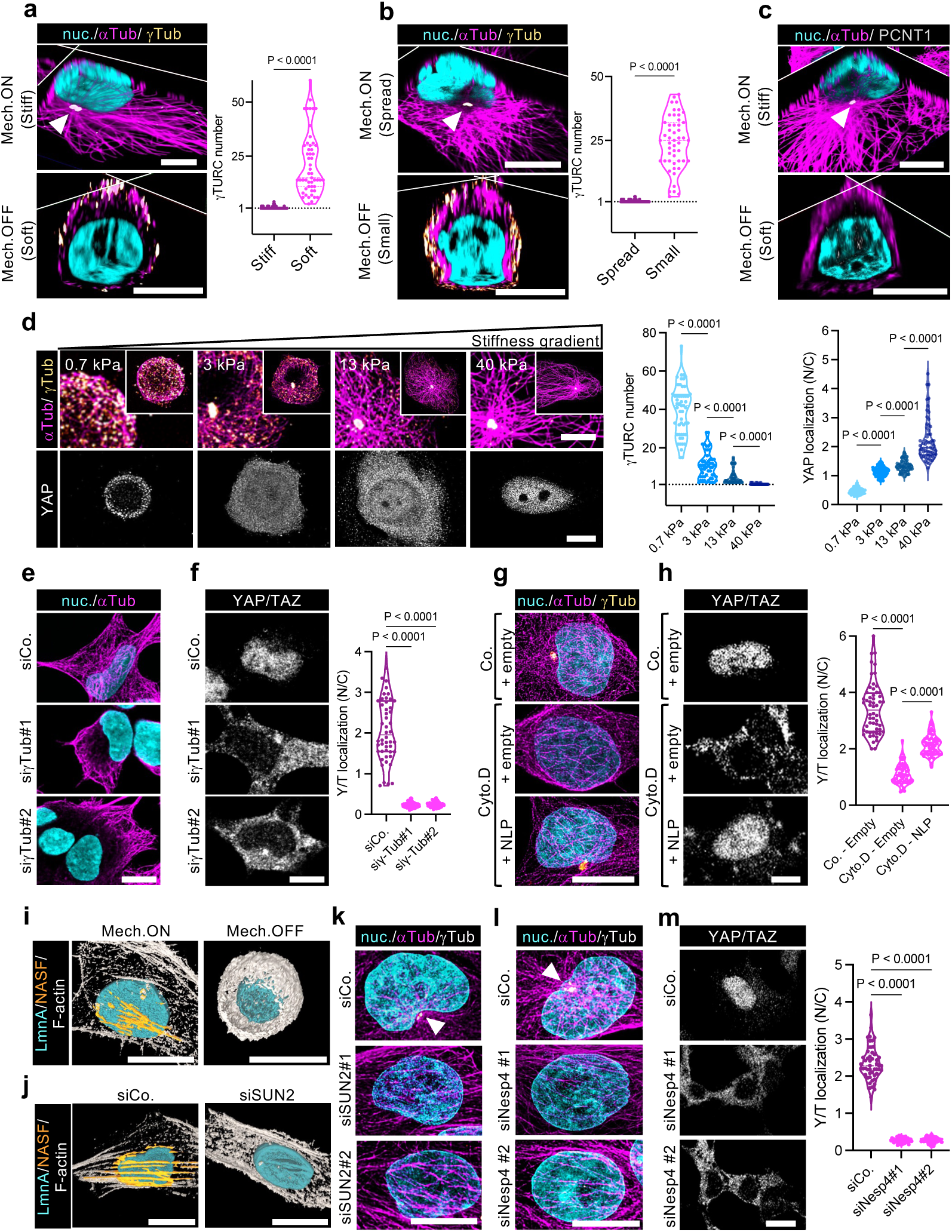
Microtubules (MTs) centrosomal organization serves as determinant of YAP/TAZ mechanotransduction. **a)** Left: representative immunofluorescence pictures (3D sections) of MCF10A cells seeded on Stiff (40 kPa, mechano-ON, hereafter Mech.ON) versus Soft (0.7 kPa, mechano-OFF, hereafter Mech.OFF) hydrogels. Right: quantifications (n≥50) of γ-TURC number in cells seeded as in left panels. **b)** Left: representative immunofluorescence pictures (3D sections) of MCF10A cells seeded on Spread (unconfined, Mech.ON) versus Small (100 μm^2^ micropatterns, Mech.OFF) substrates. Right: quantifications (n≥50) of γ-TURC number in cells seeded as in left panels. In (**a**) and (**b**) MT architecture was labelled by α-Tubulin staining (αTub, magenta), γ-TURCs were labelled by γ-Tubulin staining (γTub, yellow) and nuclei were counterstained with Hoechst (cyan). White arrowheads in Mech.ON panels indicate γ-TURCs convergence into a single MTOC. Scale bar, 5μm. See also Extended Data Fig. 1a and Supplementary Video 1 for 3D reconstructions of cells seeded as in (**a**) and (**b**). Comparable results were obtained with HEK293 and U2OS cells (Extended Data Fig. 1c,d). **c)** Representative immunofluorescence pictures (3D sections) of MCF10A cells seeded on Stiff (40 kPa, Mech.ON) versus Soft (0.7 kPa, Mech.OFF) hydrogels. MT architecture was labelled by α-Tubulin staining (αTub, magenta), the centrosome was labelled by pericentrin staining (PCNT1, grey) and highlighted by a white arrowhead. Nuclei were counterstained with Hoechst (cyan). Scale bar, 5μm. See also Extended Data Fig. 1b for representative immunofluorescence pictures (3D sections) of MCF10A cells seeded on Spread versus Small substrates and stained as in (**c**). Comparable results were obtained with HEK293 and U2OS cells (Extended Data Fig. 1c,d). **d)** Left: Representative immunofluorescence images of MTs and γ-TURCs (top panels) and EGFP-YAP (bottom panels) in MCF10A cells bearing a YAP-EGFP knock-in^(KI)^ allele seeded on hydrogels tuned to the indicated stiffness gradient. Images of the top panels are magnifications of the insets shown in the upper right corner of each picture. Scale bar 5μm. Right: quantifications (n≥50) of γ-TURCs (left) and YAP nuclear-to-cytoplasmic subcellular localization (N/C, right) in cells seeded as in left panels. **e)** Representative immunofluorescence images of MTs (αTub) in HEK293 cells transfected with the indicated siRNAs. Nuclei were counterstained with Hoechst (cyan). Scale bar, 10μm. **f)** Left: representative immunofluorescence images of YAP/TAZ (grey) in HEK293 cells transfected with the indicated siRNAs. Right: quantifications (n≥50) of YAP/TAZ nuclear-to-cytoplasmic subcellular localization (N/C) in cells treated as in left panels. **g)** Representative immunofluorescence images of MTs (αTub) and γ-TURCs (γTub) in HEK293 cells transduced with empty or NLP1-encoding lentiviruses and treated with Cytochalasin D (Cyto.D, 1 μM for 2 hrs) or DMSO as negative control (Co.). Nuclei were counterstained with Hoechst (cyan). Scale bar, 5μm. **h)** Left: representative immunofluorescence images of YAP/TAZ (grey) in HEK293 cells treated as in (**g**). Right: quantifications (n≥50) of YAP/TAZ nuclear-to-cytoplasmic subcellular localization (N/C) in cells treated as in left panels. See also Extended Data Fig. 2h for controls showing effective F-actin disruption by Cyto.D treatment. **i)** Representative 3D immunofluorescence reconstructions of MCF10A cells seeded in Mech.ON (40 kPa) versus Mech.OFF (0.7 kPa) conditions. Scale bar, 10μm. See also Supplementary Video 2 for a wrap-around view and Extended Data Fig. 3a for representative Z-planes of cells seeded as in (**i**). The nuclear lamina was labelled by LaminA staining (LmnA, cyan), F-actin was labelled by Phalloidin staining (F-actin, grey). Nuclear associated stress fibers (NASFs) are highlighted in orange in the 3D reconstruction (see methods). See also Supplementary Video 2. **j)** Representative 3D immunofluorescence reconstructions of MCF10A cells treated with the indicated siRNAs. Scale bar, 10μm. See also Extended Data Fig. 3d for representative Z-planes of cells treated with the same siRNAs. The nuclear lamina was labelled by LaminA staining (LmnA, cyan), F-actin was labelled by Phalloidin staining (F-actin, grey). Nuclear associated stress fibers (NASFs) are highlighted in orange in the 3D reconstruction (see methods). **k,l)** Representative immunofluorescence images of MTs (αTub) and γ-TURC (γTub) in HEK293 cells transfected with the indicated siRNAs. Nuclei were counterstained with Hoechst (cyan). White arrowhead indicates MTs convergence in a perinuclear MTOC only in control cells (siCo.). Scale bar, 10μm. **m)** Left: Representative immunofluorescence images of YAP/TAZ (grey) in HEK293 cells transfected with the indicated siRNAs. Scale bar, 10μm. Right: quantifications (n≥50) of YAP/TAZ nuclear-to- cytoplasmic subcellular localization (N/C) in cells treated as in left panels. *P* values were determined by unpaired Student’s *t*-test with Welch’s correction (**a,b**) or one-way ANOVA with Welch’s correction (**d,f,h,m**).

Next, we asked to what extent MT centrosomal organization correlates with mechanotransduction, as quantified by YAP/TAZ nuclear entry, an established proxy of YAP/TAZ activation ^19^. For this, we exposed cells to a gradient of mechanical forces by seeding them on substrates ranging from 0.7 kPa to 40 kPa elastic modules; in these cultures, we then quantified by immunofluorescence (IF) YAP/TAZ nucleo/cytoplasmic ratio and the number of γ-TURCs per individual cell by anti-γ-Tub staining (Fig. 1d). Cells with the highest level of nuclear YAP/TAZ were those displaying a single γ-TURC signal (set to 1 in Fig. 1d), corresponding to a single perinuclear MTOC. Conversely, cells seeded at the lowest stiffness displayed cytoplasmic YAP/TAZ accompanied by multiple independently scattered γ-TURCs that never converge into an individual MTOC (Fig. 1d). Intriguingly, at intermediate mechanical states (3-13 kPa in Fig. 1d), one MTOC typically coexists in the same cell with peripherally scattered γ-TURC signals, whose number correlated with YAP/TAZ cytoplasmic retention (proxy of YAP/TAZ turn OFF). These findings indicate that the perinuclear MTOC and its associated MT aster are mechanically regulated cellular structures connecting cellular architectural features with YAP/TAZ mechanotransduction.

Given the above correlations, we next asked to what extent centrosomal MT organization is causal for YAP/TAZ regulation. For this, we first inhibited MTOC formation in mechano-ON cells by depletion of γ-Tubulin. Notably, as shown in Fig. 1e, this treatment causes disappearance of radial MTs from the perinuclear area yet preserving shorter MTs at the cell periphery. Importantly, this invariably results in YAP/TAZ cytoplasmic retention (Fig. 1f), with ensuing inhibition of YAP/TAZ transcriptional activity, as measured by endogenous targets genes expression (Extended Data Fig. 1e), as such suggesting the key role of pericentrosomal MT organization for YAP/TAZ activation. Of note, cells depleted of γ-Tubulin display overtly normal Focal adhesions, as assessed by Paxillin staining, and F-actin arrangement (Extended Data Fig. 1f), a finding compatible with the view that MTs are essential regulators of YAP/TAZ function acting either downstream or in parallel to actomyosin.

To further validate that MT centrosomal organization serves as determinant of mechanosignaling, we next tackled MT acetylation, a central post-translational modification essential for MTs to bend without breaking, and thus critical for the stability of the long-lived aster of perinuclear MTs ^20^. We found that indeed in mechano-ON cells perinuclear astral MTs are more acetylated than peripheral ones, whereas mechano-OFF cells display a cage-like structure of curved and homogenously acetylated MTs (Extended Data Fig. 2a). The main enzyme responsible for MT acetylation is the Tubulin acetyl-transferase (α- TAT1) ^20^, and its depletion with independent siRNAs caused loss of pericentrosomal MTs and YAP/TAZ nuclear exclusion (Extended Data Fig. 2b-g). Of note, this occurred while preserving short MTs at the cell periphery and without interfering with F-actin architecture, focal adhesion engagement and cell spreading (Extended Data Fig. 2e), all in all phenocopying γ-Tubulin depletion.

Given the above requirement of the MTOC and radial MT sprouting for YAP/TAZ activation in mechano-ON cells, we then asked whether experimentally sustaining such structural features in mechano-OFF cells is sufficient to rescue nuclear YAP/TAZ accumulation. For this, we considered NLP1, a centrosomal factor instrumental for centrosome maturation and microtubule nucleation ^21^. We opted to overexpress NLP1 in cells experiencing an extreme mechano-OFF condition, that is, in cells void of F-actin after Cytochalasin D (Cyto. D) treatment. Remarkably, even in this condition, gain of NLP1 was sufficient to rescue perinuclear MTOC formation, radial MTs and YAP/TAZ nuclear accumulation (Fig. 1g,h and Extended Data Fig. 2h). As corollary, these results also imply that the centrosome acts formally downstream of F-actin, as forcing centrosome formation is sufficient to sustain mechanosignaling in mechano-OFF conditions.

### A mechanical continuum between a stress fiber subpool, the nuclear envelope and microtubules

Results above leave unaddressed how mechanical forces, well-known to be primarily perceived by F-actin at sites of cell-ECM adhesions ^8, 9^, could ultimately converge on perinuclear MT organization and YAP/TAZ regulation. The nuclear envelope (NE) has recently emerged as central hub integrating tensional forces generated by the actomyosin cytoskeleton in a mechanical continuum with the other cytoskeletal systems ^16^. Central in this network are LINC complexes, that connect nuclear Lamins with F-actin (through Nesprin1/2 and SUN) and MTs (through Nesprin4) ^16, 17, 22^. We found by dual IF for F-actin and LaminA that two distinct subpools of stress fibers exist in epithelial cells: stress fibers that do not contact the nuclear envelope, and ventral stress fibers that do (hereafter Nuclear-Associated-Stress-Fibers, NASFs) (Fig. 1i and Extended Data Fig. 3a). This distinction was derived by applying a machine-learning classifier to differentially segment phalloidin-stained F-actin fibers whose pixels colocalize with the LaminA-stained nuclear envelope in 3D confocal reconstructions. Importantly, NASFs tether the cell surface with the basal side of the NE exclusively in mechano-ON cells (see also Supplementary Video 2). Using a conformation sensitive anti-LaminA antibody ^23^, we also found that the NASFs-NE interaction stretches the basal side of the NE (Extended Data Fig. 3b). Interrupting such mechanical continuum by SUN2 depletion indeed not only abolishes NASFs formation and induces NE relaxation (Fig. 1j and Extended Data Fig. 3c,d), but also causes YAP/TAZ cytoplasmic retention with ensuing downregulation of YAP/TAZ target genes (Extended Data Fig. 3e-h). Of note, this occurs without overtly interfering with cell shape or other stress fiber subtypes that characterize the mechano-ON state (Extended Data Fig. 3d, i).

MTs are known to push against the nucleus, inducing the typical indentation described to host the centrosome in many cultured mammalian cells (“centrosomal bay”) ^24, 25^. Is NE anchorage by NASFs required for such MT-generated forces? We found that SUN2 depletion impeded close MT attachment to the NE, in turn causing defective MTOC formation and localization, all in all resulting in severely inhibited MT convergence to the perinuclear centrosome (Fig. 1k). Then we asked whether the physical connection between MTs and the NE is key to allow centrosome formation and MT radial organization. For this, we interfered with Nesprin 4 or Emerin, the main MT-anchoring factors at the NE ^16, 17, 22^. Similarly to SUN2 depletion, loss of Emerin or Nesprin 4 results in defective MTOC, loss of centrosomal MTs and YAP/TAZ inhibition (Fig. 1l,m and Extended Data Fig. 4a-h).

To reinforce the notion that MT centrosomal organization acts downstream of F-actin-mediated force sensing, we monitored YAP/TAZ activity in cells lacking the F-actin capping/severing factors ADF/Cofilin. As previously reported, depletion of ADF/Cofilin potently rescued YAP/TAZ activity in mechano-OFF conditions ^10^. However, we found that this rescue is impaired by concomitant depletion of α-TAT1 or Nesprin4 (Extended Data Fig. 4i), indicating that boosting F-actin stress fibers requires pericentrosomal MTs to impact on mechanosignaling.

#### AMOT protein stabilization as mechano-rheostat

How does the centrosomal MTOC trigger YAP/TAZ mechanotransduction? In addressing this crucial question, we considered the well-known role of the multiprotein centrosomal assembly in coordinating local degradation of proteasomal substrates. Indeed, prior work has shown that other signaling factors - such as polyubiquitinated Smad1, β-catenin and Axin2 - display continuous delivery to the centrosomal proteasome ^26–28^. Also well-established is the role of MTs originated from the MTOC in allowing dynein-mediated retrograde flow of cellular components toward the proteasome at pericentrosomal condensates ^15^. As previously reported ^27, 29^, the cellular proteasome is enriched at the centrosomal bay (Fig. 2a). We found that proteasomal accumulation at the centrosome itself is dependent on the mechanical state of the cell and correlates with MTs architecture, with mechano-ON cells displaying proteasome enrichment at the perinuclear centrosome and mechano-OFF cells displaying more diffused proteasomes throughout the cytoplasm (Fig. 2a and Extended Data Fig. 5a). Intriguingly, we also found that proteasome localization is dependent on the NASFs-NE-MTs mechanical equilibrium, as interfering with SUN2 or Emerin results in failed proteasome aggregation at the centrosomal bay (Fig. 2b and Extended Data Fig. 5b).

**Fig. 2:**
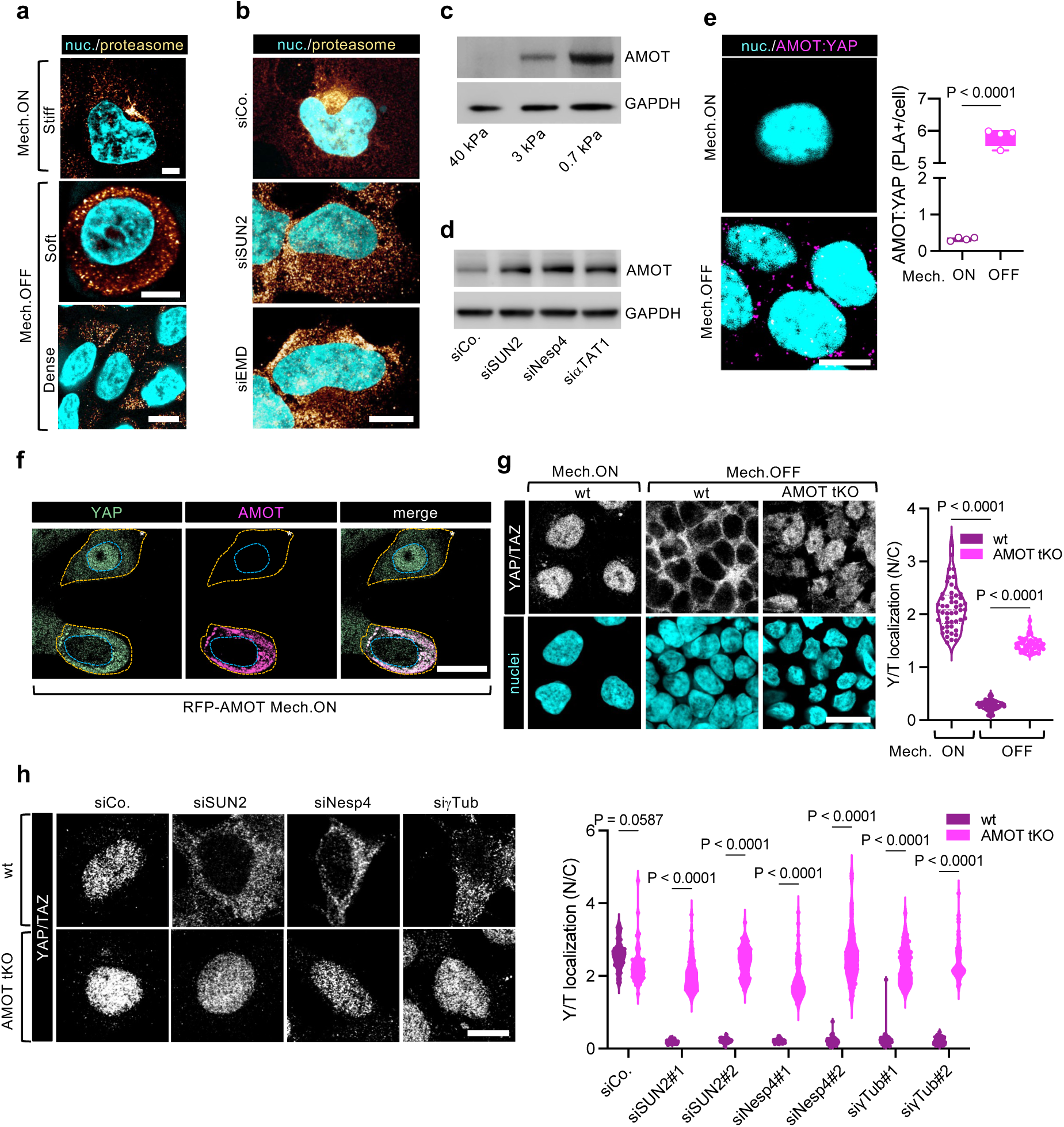
AMOT acts as cytoplasmic mechanical rheostat to control YAP/TAZ activity. **a)** Representative immunofluorescence images of HEK293 cells seeded in mechano-ON (Stiff, 40 kPa hydrogels) versus mechano-OFF (Soft, 0.7 kPa hydrogels or dense culture) conditions. The proteasome was labelled by 20S/PSMA5 staining (proteasome, yellow) and nuclei were counterstained with Hoechst (cyan). Scale bar, 10μm. See also Extended Data Fig. 5a for quantifications of proteasome localization in cells seeded in the same conditions. **b)** Representative immunofluorescence images of HEK293 cells seeded in Mech.ON conditions and treated with the indicated siRNAs. The proteasome was labelled by 20S/PSMA5 staining (proteasome, yellow) and nuclei were counterstained with Hoechst (cyan). Scale bar, 10μm. See also Extended Data Fig. 5b for quantifications of proteasome localization in cells treated in the same way. **c)** Representative AMOT immunoblot of MCF10A cells seeded on hydrogels of the indicated stiffness. GAPDH serves as loading control. See also Extended Data Fig. 5d,f-h for AMOT immunoblots from cells experiencing independent mechano-OFF conditions by treatment with F-actin inhibitors, and Supplementary Fig. 1a for quantifications. **d)** Representative AMOT immunoblot of HEK293 cells treated with the indicated siRNAs. GAPDH serves as loading control. See also Supplementary Fig. 1a for quantifications. **e)** Left: representative images of proximity ligation assays (PLA) showing exclusively cytoplasmic interaction (magenta) between endogenous AMOT and YAP/TAZ in HEK293 cells seeded in Mech.OFF conditions. Mech.ON conditions (in absence of endogenous AMOT protein) serve as negative control. Nuclei were counterstained with Hoechst (cyan). Scale bar, 10μm. Right: quantifications of the mean number of PLA dots per cell (>50 cells were quantified for each independent experiment, indicated as dot, n=4). **f)** Representative stills from live fluorescence images of RFP-AMOT expressing MCF10A-YAP-EGFP^KI^ cells showing that AMOT overexpression is sufficient to cause YAP cytoplasmic retention in mechano- ON conditions. Cells without AMOT overexpression (*), showing nuclear YAP accumulation, serve as negative control. Cell and nuclear borders are outlined in yellow and blue dashes, respectively. Scale bar, 20μm. **g)** Left: representative immunofluorescence images of YAP/TAZ in control (wt) versus AMOT- 130/AMOT-L1/AMOT-L2 triple KO (AMOT tKO) HEK293 cells in Mech.ON vs Mech.OFF conditions. Nuclei were counterstained with Hoechst (cyan). Scale bar, 20μm. Right: quantifications (n≥50) of YAP/TAZ nuclear-to-cytoplasmic subcellular localization (N/C) in cells seeded as in top panels. **h)** AMOT is epistatic to LINC proteins and MTs. Left: representative immunofluorescence images of YAP/TAZ in control versus AMOT tKO HEK293 cells treated with the indicated siRNAs. Scale bar, 10μm. Right: quantifications (n≥45) of YAP/TAZ nuclear-to-cytoplasmic subcellular localization (N/C) in cells treated as in top panels. *P* values were determined by unpaired Student’s *t*-test with Welch’s correction (**e**) or one-way ANOVA with Welch’s correction (**g,h**).

We surmised that a scenario that could mechanistically explain YAP/TAZ regulation by MTs may be one in which the mechanoregulated perinuclear MTOC and proteasome condensation are required for degradation of a YAP/TAZ cytoplasmic inhibitor. Although the existence of a mechano-regulated YAP/TAZ inhibitor has been long postulated, its identification has remained so far elusive ^30^. In pursuing this hypothesis, we unbiasedly searched the BioGRID4.4 dataset ^31^ to compile a list of cytoplasmic proteins interacting with both YAP and TAZ (Table 1), and tested whether mechanical cues could tune their endogenous levels. By carrying out western blots of mechano-ON vs. -OFF cell lysates, we found that the stability of only one of such proteins, AMOT-p130, stood out for being strikingly mechano- regulated (Fig. 2c and Extended Data Fig. 5c,d), in absence of any effect at the level of AMOT mRNA transcription (Extended Data Fig. 5e). This was an intriguing finding as AMOT has been shown to interact directly with YAP/TAZ ^32^ acting as a scaffold protein for their LATS-mediated phosphorylation, as such contributing to YAP/TAZ inhibition by the Hippo pathway ^33^. Yet, the role of AMOT in mechanosignaling remains largely unaddressed. In vitro, AMOT has also been proposed to associate with F-actin ^34^, although the significance of this observation for mechanosignaling remains unclear. Indeed, we found that AMOT levels are equally stabilized by either treatments with microfilament inhibitory drugs (Extended Data Fig. 5f-h) or physiologically mimicking mechano-OFF cell states (Fig. 2c), in which F-actin levels, if anything, tend to increase when compared to mechano-ON cells (Extended Data Fig. 5i). Importantly, we also found that AMOT levels are also similarly increased by treatment of mechano-ON cells with inhibitors if microtubule polymerization, such as Nocodazole and Vincristine (Extended Data Fig. 5j). In other words, AMOT stabilization is orchestrated by mechanosignaling events formally operating downstream of F-actin.

Next, we thus addressed the role of centrosomal MT organization as determinant of AMOT stabilization. We tested this in all experimental conditions that we found above to inhibit centrosome formation. We found that AMOT stabilization does occur in mechano-ON cells after: i) uncoupling the NE from NASFs (i.e., after SUN2 depletion); ii) uncoupling the NE from MTs (i.e., upon Nesprin4 depletion); or iii) impairing perinuclear MT aster stability through inhibition of MT acetylation (i.e., after α-TAT1 depletion) (Fig. 2d).

Is AMOT stability a mechanical rheostat responding to MT organization for YAP/TAZ regulation? To address this, we used proximity ligation assays to spatially map YAP/AMOT interaction and found that this occurs exclusively in the cytoplasm of mechano-OFF cells (Fig. 2e). Moreover, supporting AMOT levels in mechano-ON cells by an AMOT expression plasmid is sufficient to quantitatively retain YAP in the cytoplasm (Fig. 2f) and to blunt YAP/TAZ transcriptional responses (Extended Data Fig. 5k,l). YAP cytoplasmic retention relies on its physical association with AMOT as demonstrated by two complementary experiments involving mutations in the known interacting domains of the two proteins: first, expression in mechano-ON cells of AMOT versions bearing point mutations that impair YAP/TAZ binding ^35, 36^ was inconsequential for YAP/TAZ activity (Extended Data Fig. 5k); second, reconstitution of YAP/TAZ-depleted cells with a YAP-WW-mutant rendered them insensitive to AMOT add-back (Extended Data Fig. 5l). Collectively, the data indicate that AMOT protein stability operates as mechanical rheostat through YAP/TAZ.

Beyond gain-of-function, we next investigated to what extent YAP/TAZ nuclear entry requires AMOT degradation. For this, we depleted all AMOT proteins (AMOTp130, AMOTL1 and L2) from mechano- OFF cells by triple CRISPR-mediated knockout (KO) or siRNAs. Remarkably, AMOT depletion fully rescued YAP/TAZ nuclear entry and activity in mechano-OFF cells (Fig. 2g). We proved this in several experimental conditions, such as in cells seeded on soft substrates or small adhesive islands (Extended Data Fig. 5m,n), in cells with impaired F-actin contractility (Extended Data Fig. 5o), and also in cells carrying centrosomal MTs impairment through depletion of SUN2, Nesprin4 or γ-Tubulin (Fig. 2h). In all these mechano-inhibitory conditions, AMOT was the downstream factor causing YAP/TAZ inhibition, as its depletion rescued YAP/TAZ nuclear localization and transcriptional activity, as measured by YAP/TAZ endogenous target genes and the YAP/TAZ/TEAD-activity synthetic reporter 8XGTIIC-Lux (Fig. 2g,h and Extended Data Fig. 5m-o).

#### Mechanical regulation of AMOT stability relies on retrograde transport along MTs

We have shown above that AMOT degradation in mechano-ON cells relies on the centrosome and MT radial organization, raising questions on the underlying mechanism. Proteasomal degradation of AMOT has been previously described downstream of poly-ADP ribosylation (i.e., “PARylation”) by TNKS1/2 enzymes serving as degron-tag to promote AMOT poly-ubiquitilation by the E3 ligase RNF146 ^37^. Consistently, as previously reported and here confirmed, inhibiting PARylation or depleting cells of the E3 ligase RNF146 potently supports AMOT stabilization (Extended Data Fig. 6a,b).

Given this cascade, we asked whether any of these biochemical events is in fact regulated by cell mechanics. Tankyrase1/2 -dependent AMOT PARylation and subsequent poly-ubiquitination occur in both mechano-ON and mechano-OFF conditions and, consistently, RNF146 protein levels and subcellular localization were also indistinguishable in mechano-ON and -OFF conditions (Extended Data Fig. 6c-f). This prompted us to consider the role of the mechano-regulated centrosomal proteasome as determinant of AMOT stability. To test this notion, we first confirmed that AMOT protein levels in mechano-ON cells could be rescued, dose dependently, by treatment with proteasome inhibitors (Fig. 3a). Next, we followed AMOT subcellular localization by 3D imaging in proteasome-inhibited mechano- ON cells. Notably, in these conditions, the sole AMOT protein that could be identified is indeed concentrated at centrosomal condensates (Fig. 3b). Consistently, we found that AMOT protein could be immunoprecipitated in complexes with the proteasome component 20S, and exclusively in cells seeded in mechano-ON conditions (Fig. 3c). Thus, in mechano-ON cells, AMOT is constantly targeted to the pericentrosomal area allowing robust, quantitative and timely control of protein levels, essential for stringent cellular mechano-responsiveness. To quantitatively probe the rate of AMOT protein degradation in mechano-ON cells, we conducted cycloheximide pulse-and-chase experiments, showing that the entire AMOT protein pool that is stabilized by proteasome inhibitor treatment in mechano-ON cells, is readily degraded after proteasome inhibitor washout (Extended Data Fig. 6h).

**Fig. 3:**
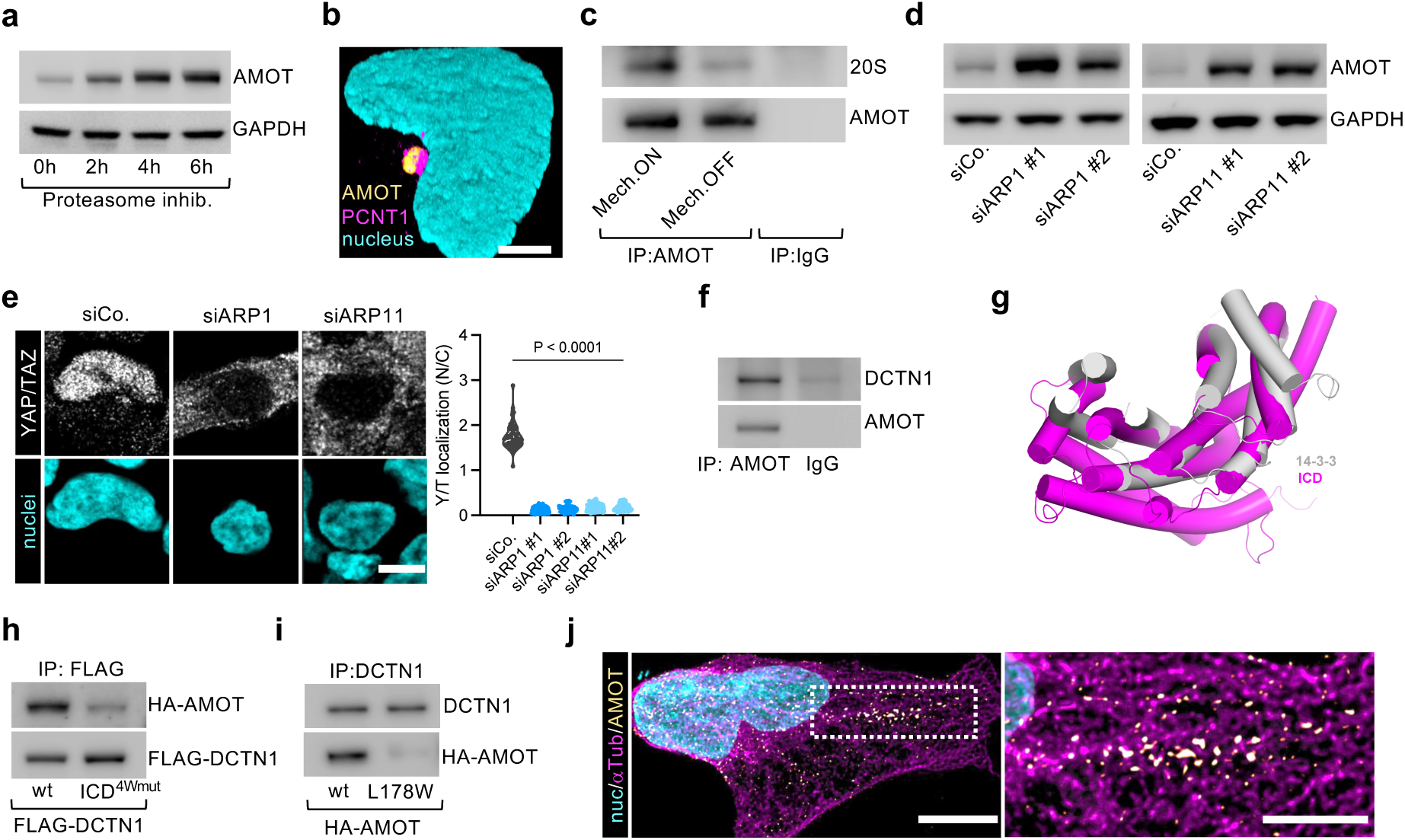
Mechanical regulation of AMOT stability relies on retrograde transport along MTs in mechano-ON cells. **a)** Representative AMOT immunoblot of HEK293 cells seeded in mechano-ON conditions and treated with proteasome inhibitor (Lactacystin 10μM) for the indicated timings. GAPDH serves as loading control. **b)** Representative 3D immunofluorescence reconstructions of HEK293 cells seeded in mechano-ON conditions treated with proteasome inhibitor (Lactacystin 10μM, 6hrs). Centrosome was labelled by pericentrin staining (PCNT1, magenta), AMOT was labelled in yellow and the nucleus was counterstained with Hoechst (cyan). Scale bar, 5μm. **c)** Pulldown of endogenous AMOT from proteasome-inhibited HEK293T cell seeded in Mech.ON versus Mech.OFF conditions, showing AMOT interaction with the proteasome component 20S exclusively in Mech.ON conditions. IgG pulldown serves as negative control. The inputs of the same pulldown experiment are shown in Extended Data Fig. 6g. **d)** Representative AMOT immunoblots of HEK293 cells treated with the indicated siRNAs targeting components of the Dynactin complex. GAPDH serves as loading control. **e)** Left: Representative immunofluorescence images of YAP/TAZ in HEK293 cells transfected with the indicated siRNAs. Nuclei were counterstained with Hoechst (cyan). Scale bar, 10μm. Right: quantifications (n≥50) of YAP/TAZ nuclear-to-cytoplasmic subcellular localization (N/C) in cells treated as in left panels. **f)** Pulldown of endogenous AMOT from proteasome-inhibited mechano-ON HEK293T cell lysates showing AMOT interaction with the Dynactin complex component DCTN1. IgG pulldown serves as negative control. **g)** Structural superposition between the 14-3-3 domain (grey) and intra-coiled-coil domain (ICD) of DCTN1/p150 (magenta). See also Extended Data Fig. 7j for predicted alignment error (PAE) plot for the complex shown here. **h)** Pulldown of FLAG-DCTN1 from mechano-ON HEK293T cells transiently transfected with HA- AMOT and the indicated FLAG-DCTN1 mutants and treated with Dynarrestin 10μM for 6hrs, showing impaired interaction between AMOT and a DCTN1 mutant bearing S659W/L662W/T835W/A839W substitutions in the ICD (ICD^4W^ mutant). The inputs of the same pulldown experiment are shown in Extended Data Fig. 7m. **i)** Pulldown of endogenous DCTN1 from mechano-ON HEK293T cells transiently transfected with the indicated AMOT mutants and treated with Dynarrestin 10μM for 6hrs, showing impaired interaction between DCTN1 and AMOTL178W mutant. The inputs of the same pulldown experiment are shown in Extended Data Fig. 7n. **j)** Representative immunofluorescence image of MTs (αTub, magenta) and endogenous AMOT (yellow) in MCF10A cells seeded in Mechano-ON conditions and treated with Dynarrestin 10μM for 6hrs. Nuclei were counterstained with Hoechst (cyan). Scalebar, 10μm. The right panel shows a magnification of the area dashed in the left panel. Scalebar, 5μm. See also Supplementary Video 3 for live time lapse videos showing AMOT retrograde transport along MTs in DMSO versus Dynarrestin-treated MCF10A cells. *P* values were determined by one-way ANOVA with Welch’s correction (**f**).

Next, we asked by what means AMOT is delivered to centrosomal condensates. Interesting clues came by considering publicly available AMOT interaction datasets. We first considered recent proteome-wide reports mapping proteins that, in living cells, colocalize within the same subcellular compartment ^38^. There, we retrieved AMOT exclusively in association to Hippo components, MTs and centrosome components (Extended Data Fig. 7a and Supplementary Table 1). We also considered published protein-protein interaction datasets ^31^ that suggested that all AMOT family members can form complexes with components of the dynein system (Supplementary Table 2), well known to mediate retrograde transport toward the MTOC.

Given these clues, we thus reasoned that AMOT might reach the centrosomal location through the same mechanism shared by most pericentrosomal factors, that is, by dynein-dependent retrograde transport along the MT conveyor belt ^15^. We first approached this model from a functional perspective, asking whether interference with dynein motor activity could impact on AMOT stability. For this, we inhibited dynein-mediated retrograde transport by depleting components of the Dynein system, that is made by two portions: the MT-binding dynein motor complex, and the Dynactin complex, that is the cargo-binding cofactor of this transport system ^39^. As shown in Fig. 3d and Extended Data Fig. 7b, depletion of Dynein-light chain 2, as well as of the Dynactin components ARP1A/B or ARP11, readily results in aberrant AMOT stabilization in mechano-ON cells. Reinforcing this conclusion, a similar effect was obtained by blocking dynein-mediated retrograde transport with Dynarrestin, an inhibitor of Dynein association to MTs (Extended Data Fig. 7c), or with Ciliobrevin D, a reversible inhibitor of Dynein ATPase activity (Extended Data Fig. 7d). Importantly, AMOT stabilization is dynamically and quantitatively regulated by dynein dependent retrograde transport towards the pericentrosomal proteasome, as AMOT is readily degraded within 4 hours after Ciliobrevin D washout (Extended Data Fig. 7d). Interfering with retrograde transport by depletion of Dynein or Dynactin components is indeed sufficient to inhibit YAP/TAZ responses (Fig. 3e and Extended Data Fig. 7e-g), as expected by the requirement of retrograde transport to blunt AMOT levels in mechano-ON cells.

The above results prompted us to test a biochemical interaction between AMOT and components of the cargo-binding Dynactin complex. The latter is in turn made by two multiprotein portions, an actin-like ARP1/11 filament binding to Dyneins, and a projecting DCTN1 “shoulder complex”, required for recruiting specific cargos ^40^. Using cell lysates of mechano-ON cells pretreated with proteasome inhibitors, we found that pull-down of endogenous AMOT specifically co-immunoprecipitated with endogenous DCTN1, but not ARP11 or ARP1 (Fig. 3f). Of note, when bound to DCTN1, AMOT is ostensibly unable to bind to YAP, as we observed interaction between endogenous AMOT and YAP exclusively in mechano-OFF conditions (Extended Data Fig. 7h).

Next, we wished to gain insights into of the interaction between AMOT and DCTN1. As revealed by electron microscopy studies ^40–42^, DCTN1 is composed of multiple coiled-coil regions with an N-terminal CAP-Gly domain responsible, together with the basic amino acid-rich region, for MTs binding and a central intra-coiled-coil domain (ICD), whose functional role is still poorly understood. The structure of an AMOT peptide has been resolved by X-ray crystallography in complex with the 14-3-3 scaffold protein, an association requiring AMOT phosphorylation in pSer175 ^43^ (Extended Data Fig. 7i). AlphaFold2 ^44, 45^ predicted with very high confidence the structure of the ICD, revealing a structural similarity between the ICD predicted structure and 14-3-3, particularly striking for the structural arrangement of the helices contributing to the concave surface by which 14-3-3 associates to AMOT (Fig. 3g and Extended Data Fig. 7j,k). The structural similarity between 14-3-3 and the ICD promoted the hypothesis that AMOT might associate to the ICD using the same region centered around S175. To address this idea, we performed co-IP experiments and found that indeed replacement of S659, L662, T835, A839 located on concave surface of the ICD (Extended Data Fig. 7k,l) with tryptophan residues impaired binding of DCTN1 to wild-type AMOT (Fig. 3h and Extended Data Fig. 7m). Conversely, the L178W AMOT variant, predicted to impair ICD association, also abolished the interaction with endogenous DCTN1 (Fig. 3i and Extended Data Fig. 7n).

Collectively, the above biochemical results support a model in which the mechanical regulation of AMOT abundance relies on its loading onto Dynein-Dynactin complexes and its constant retrograde delivery to the proteasomal sink along the pericentrosomal MT aster. Supporting this model, by co-staining AMOT and MTs in mechano-ON cells treated with Dynarrestin, we found that AMOT forms discrete condensates decorating MT tracks and is enriched in the pericentrosomal area (Fig. 3j). Moreover, following by live imaging the movement of AMOT condensates on MTs, we were able to directly visualize a very fast retrograde transport of RFP-tagged AMOT aggregates along MTs, a flow that is dramatically reduced by dynarrestin treatment (Supplementary Video 3).

#### A unifying model integrating mechanosignaling and the Hippo cascade

The above results offer fresh perspectives on a currently debated issue, that is how the Hippo pathway intersects with mechanosignaling. Prior work has shown that AMOT is phosphorylated by the core Hippo kinases LATS1/2 ^32, 46, 47^. Here we found that LATS1/2 activity contributes to AMOT stability, as demonstrated by AMOT enhanced degradation in LATS1/2 depleted cells (Fig. 4a and Extended Data Fig. 8a). What attracted our attention was that LATS1/2 phosphorylates AMOT-130 exactly in S175 (Extended Data Fig. 8b), namely, within the same region responsible for loading AMOT on 14-3-3 ^43^ and on the DCTN1/dynactin complex (see above Fig. 3g and Extended Data Fig. 7i). This raised the possibility that AMOT phosphorylation by LATS may tip the balance of these associations averting AMOT from Dynactin. To test this, we used HA-tagged AMOT phospho-mimetic AMOT-S175E and phospho-mutant AMOT-S175A constructs and carried out reciprocal co-IP/western blot experiments to monitor their association with endogenous DCTN1 in cell lysates. We found that AMOT-S175E and AMOT-S175A showed lower and higher ability to bind to DCTN1, respectively (Fig. 4b,c and Extended Data Fig. 8c,d). Consistently, after dose titration and western blotting, we found that AMOT-S175E is a more stable protein compared to AMOTwt, whereas AMOT-S175A is readily degraded (Extended Data Fig. 8e). In sum, LATS-AMOT phosphorylation represents a “salvage” mechanism favoring AMOT escape from its degradation pathway.

**Fig. 4:**
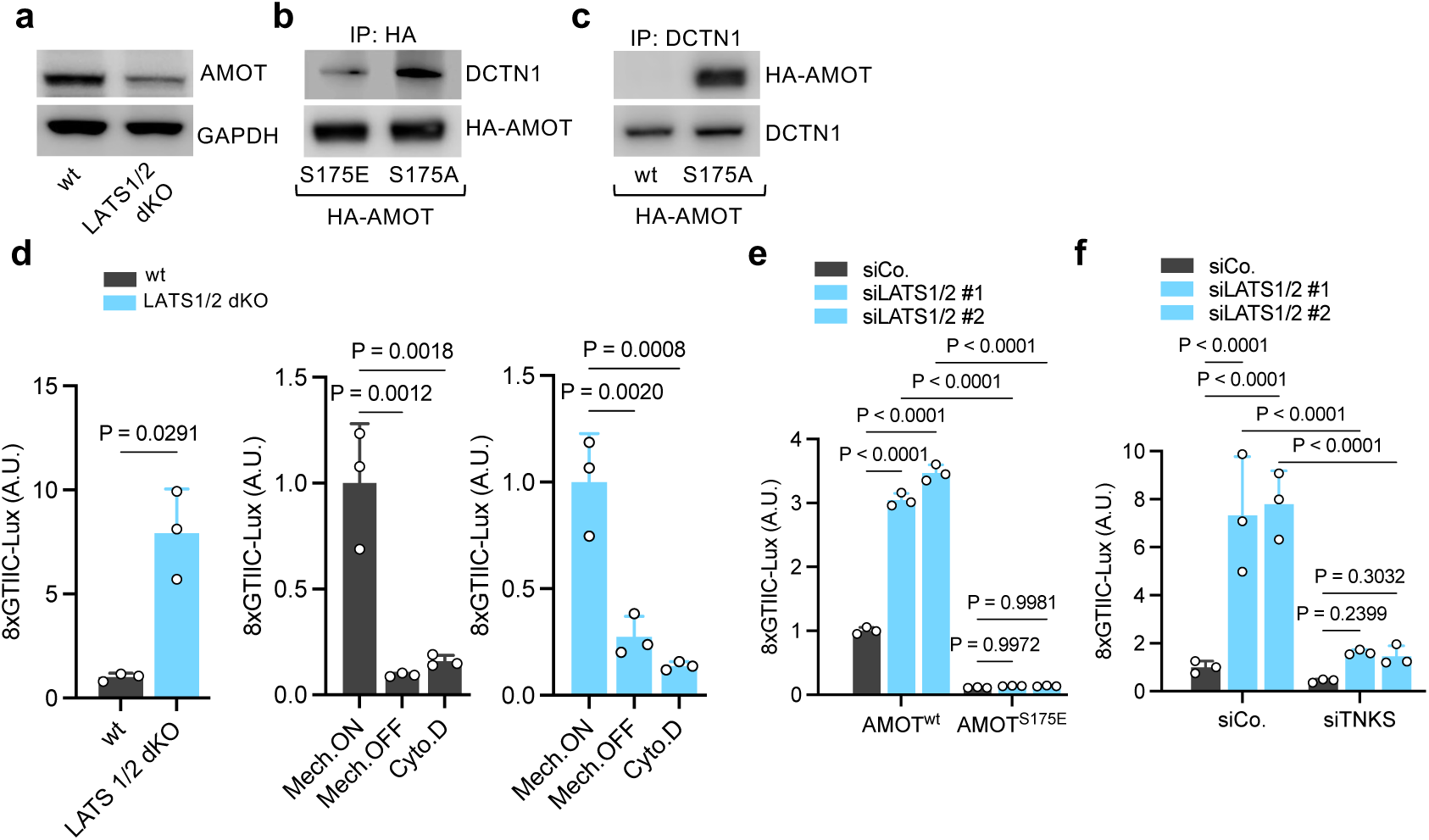
Hippo/LATS signaling regulates YAP/TAZ mechanotransduction through AMOT. **a)** Representative immunoblots of mechano-OFF HEK293 LATS1/2 dKO cells showing impaired AMOT stabilization upon LATS1/2 double knockout. GAPDH serves as loading control. **b)** Pulldown of HA-AMOT from mechano-ON HEK293T cells transiently transfected with the indicated HA-AMOT mutants and treated with Lactacystin 10μM for 6hrs, showing lower and higher ability of the AMOT-S175E and AMOT-S175A mutants to bind to DCTN1, respectively. The inputs of the same pulldown experiment are shown in Extended Data Fig. 8c. **c)** Pulldown of endogenous DCTN1 from mechano-ON HEK293T cells transiently transfected with the indicated HA-AMOT mutants and treated with Dynarrestin 10μM for 6hrs, showing enhanced interaction between DCTN1 and AMOT S175A mutant. The inputs of the same pulldown experiment are shown in Extended Data Fig. 8d. **d)** Luciferase assay of control (wt) or LATS1/2 dKO HEK293 cells transfected with a synthetic reporter for YAP/TAZ/TEAD-dependent transcription (8xGTIIC-Lux). Middle and right panels show luciferase assays performed with these same cells seeded in Mech.ON (sparse), Mech.OFF (dense) conditions or treated with Cyto.D (0.5 μM, 15 hours). Data are mean + s.d. of n = 3 biologically independent samples. **e)** Luciferase assay of HEK293 cells treated with the indicated siRNAs and transfected with a synthetic reporter for YAP/TAZ/TEAD-dependent transcription (8xGTIIC-Lux) and with the indicated AMOT mutants. Data are mean + s.d. of n = 3 biologically independent samples. **f)** Luciferase assay of HEK293 cells treated with the indicated siRNAs and transfected with a synthetic reporter for YAP/TAZ/TEAD-dependent transcription (8xGTIIC-Lux). Data are mean + s.d. of n = 3 biologically independent samples. *P* values were determined by unpaired Student’s *t*-test with Welch’s correction (**d**, left panel), one-way ANOVA with Tukey’s multiple comparison test (**d**, middle and right panels), or two-way ANOVA (**e,f)**.

The above results depict a scenario in which AMOT serves as central rheostat of mechanical cues, also integrating key inputs from Hippo/LATS1/2 activity. Of note, by monitoring LATS1/2 activity, we also found that the levels of LATS1 autophosphorylation (a hallmark of LATS activation ^48^), and the very same AMOT phosphorylation by LATS, are ostensibly indistinguishable in lysates from mechano-ON and -OFF cells (Extended Data Fig. 8f). In other words, this suggests that AMOT regulation by LATS is a tonic modulation that constantly attempts to skew AMOT from degradation to stabilization, without affecting mechano-responsiveness per se but rather modulating its amplitude. In line, LATS1/2 double knockout cells display enhanced YAP/TAZ activity but are still readily inhibited by mechano-OFF conditions or cytoskeletal inhibition (Fig. 4d).

If LATS activity in fact converges on YAP/TAZ mechanoregulation by ultimately regulating AMOT protein levels, then we should expect YAP/TAZ inhibition by AMOT to be formally uncoupled from LATS activity. To test this, we monitored YAP/TAZ activity in LATS1/2-depleted cells upon AMOT reconstitution. As shown in Fig. 4e, reconstitution with AMOT phosphomimicking (S175E) mutant is sufficient to render YAP/TAZ activity insensitive to LATS1/2 depletion, implying that the effect of LATS1/2 depletion on YAP/TAZ activity is, to a substantial extent, due to modulation of AMOT degradation. If LATS operates on YAP/TAZ mechanosignaling indirectly, by saving AMOT from its degradation pathway, then stabilizing endogenous AMOT by alternative means should render LATS dispensable for YAP/TAZ inhibition. We confirmed this prediction by promoting stabilization of endogenous AMOT by inhibiting its upstream degradation signal (i.e. by depletion of TNKS1/2), finding that this renders YAP/TAZ insensitive to inhibition by LATS1/2 (Fig. 4f).

The results above in LATS1/2 depleted cells also imply at least in part a departure from the classic modality by which the Hippo pathway has been envisioned, that is by LATS1/2-mediated direct phosphorylation of YAP/TAZ. Indeed, in mechano-OFF conditions displaying increased AMOT levels, YAP/TAZ phosphorylation *per se* should be dispensable for their inhibition. In line, cells expressing LATS-insensitive mutant isoforms of YAP could still be readily inhibited by mechano-OFF conditions (Extended Data Fig. 8g), also consistently with prior reports ^5, 10, 49–59^. In other words, experimentally uncoupling AMOT from its degradation pathway renders LATS1/2 substantially irrelevant for mechano-dependent YAP/TAZ modulation, implying that the effect of LATS1/2 depletion on YAP/TAZ activity is largely due to modulation of AMOT degradation (see schematic representations of Fig. 6a,b).

#### Biological role of AMOT in classic mechanobiology assays

The data above imply a central functional role of AMOT for biological responses driven by YAP/TAZ mechanical regulation. We thus set to validate this notion in classic mechanobiology assays. Mesenchymal stem cell differentiation toward an adipogenic fate is known to be strictly dependent on YAP/TAZ mechanical inhibition ^5^. We found that AMOT is crucial in this context, as AMOT depletion impairs adipogenic differentiation in mechano-OFF states (Fig. 5a and Extended Data Fig. 9a,b); conversely, AMOT add-back is sufficient to drive adipogenic differentiation in mechano-ON mesenchymal stem cells, in which this fate is otherwise inhibited (Extended Data Fig. 9c).

**Fig. 5:**
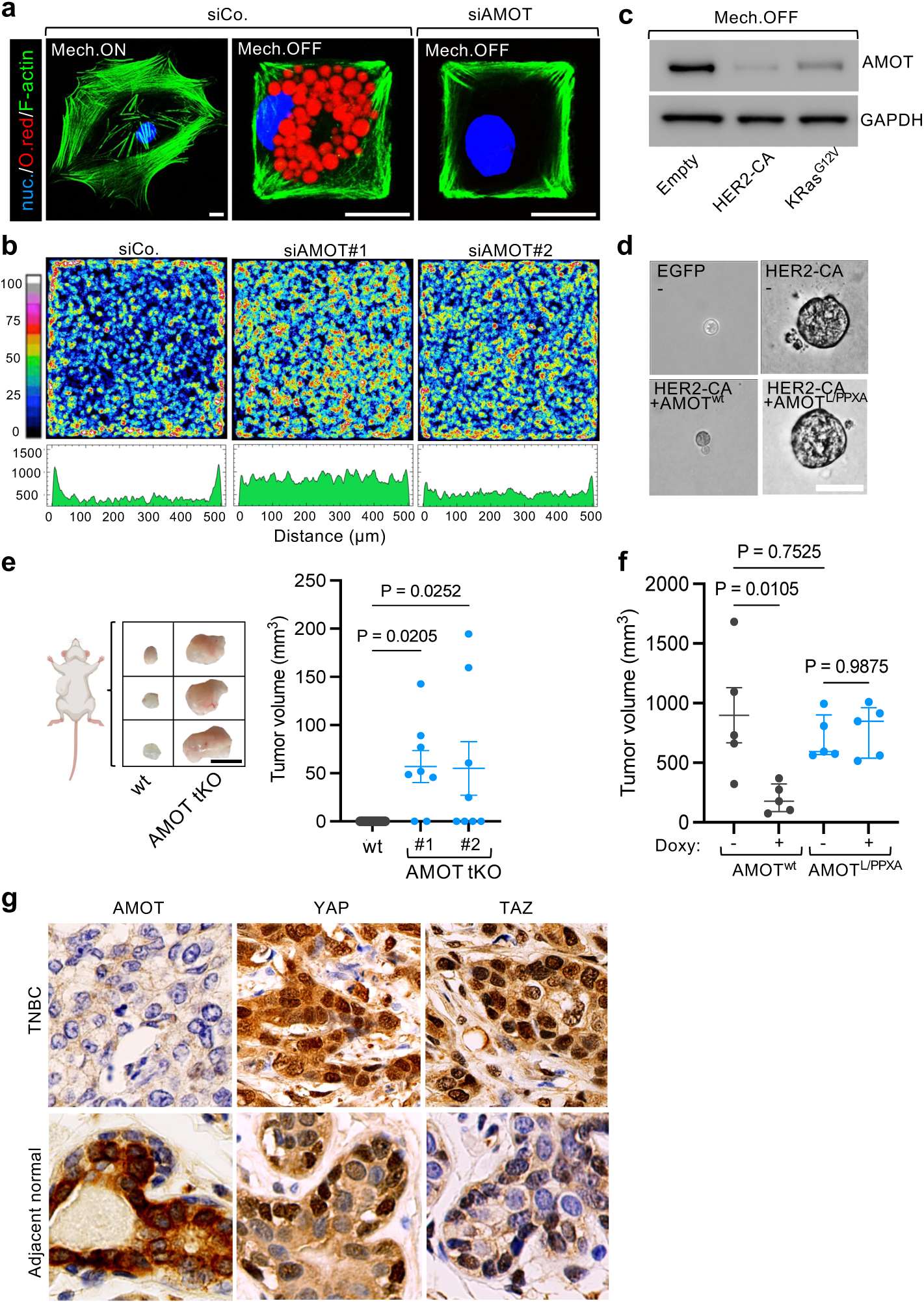
AMOT plays a central role in biological responses driven by YAP/TAZ mechanical regulation. **a)** Representative immunofluorescence pictures of hMSCs seeded on spread (unconfined, Mech.ON) or small (1000μm^2^, Mech.OFF) micropatterns and treated with the indicated siRNAs. Oil-red (O.red) staining was used to identify lipid vacuoles (red), F-actin was stained with Phalloidin (green) and nuclei were counterstained with Hoechst (blue). Scale bars, 15μm. See also Extended Data Fig. 9a for quantifications of the percentage of adipogenic differentiation in the same conditions. **b)** Top: panels show colorimetric stacked images of BrdU incorporation, used to visualize proliferation patterns of MCF10A treated with the indicated siRNAs and seeded as cell monolayers of defined dimensions. The color scale indicates the extent of cell proliferation in a given position of the monolayers. Bottom: spatial mapping of the proliferation rate (BrdU signal) along the X axis of each panel shown on top (see methods). See also Extended Data Fig. 9d for controls showing comparable density of cells in all conditions. **c)** Representative AMOT immunoblots of HEK293 cells transiently transfected with empty vector or with constitutively active oncogenes and seeded in mechano-OFF conditions. GAPDH serves as loading control. **d)** Representative images and quantifications of colonies formed by human mammary luminal differentiated cells undergoing oncogenic reprograming when transduced with lentiviral vectors encoding for the indicated factors. HER-CA, constitutive active HER2 mutant (see methods); AMOT^L/PPXA^, AMOT point mutant in the three L/PPXY YAP-binding motifs (see methods). Scale bars, 170μm. See also Extended Data Fig. 9e for quantifications. **e)** Representative images (left) and tumor volume quantifications (right, n≥8 mice for each condition) of orthotopic mammary tumors formed by control or AMOT tKO MII cells. **f)** Tumor volume quantifications (n=5 mice for each condition) of orthotopic mammary tumors formed by MDA-MB-231 cells transduced with lentiviral vectors encoding for the indicated doxycycline (Doxy)-dependent AMOT mutants. **g)** Representative immunohistochemical pictures of AMOT, YAP and TAZ proteins in chemo-naïve human triple negative breast cancer (TNBC) samples or adjacent normal mammary tissue. Data are representative of n=7 independent patient samples. See also Extended Data Fig. 10a for lower magnifications of the same samples and Extended Data Fig. 10b for quantifications of immunohistochemical staining in all tested samples. *P* values were determined by one-way ANOVA with Tukey’s multiple comparison test (**e.f**).

Next, we asked whether YAP/TAZ inactivation by AMOT dictates proliferation patterns of epithelial monolayers instructed by geometrical distribution of mechanical signals. To test this, we employed microfabricated FN islands of defined area ^10, 60^. In this setting, cells stretched at the border of the island are endowed with YAP/TAZ-dependent proliferation capacity, whereas cells at the center of the island experience density-dependent YAP/TAZ turn off and contact-inhibition of proliferation ^10^. Notably, we found that proliferation and YAP/TAZ patterning is in fact driven by AMOT patterning, with AMOT depletion resulting in aberrant proliferation throughout the entire island (Fig. 5b and Extended Data Fig. 9d).

#### AMOT as mechanoregulated tumor suppressor

Mammary tumorigenesis has long been associated to an increased tissue stiffness, and, consistently, several studies have proven that YAP/TAZ activation is indeed chiefly involved in mammary cell transformation and tumor growth ^4^. Given our results above, YAP/TAZ-dependent tumorigenesis should be expected to be favored by destabilization of AMOT. We tested this notion in several models. Tumorigenic reprogramming of mammary luminal differentiated cells by activated oncogenes, such as overexpression of constitutive active HER2, was previously shown to require mechanical forces in a YAP/TAZ-dependent manner ^49, 61, 62^. Here we found that such oncogene-mediated reprogramming is indeed accompanied by oncogene-induced AMOT degradation (Fig. 5c). Indeed, oncogene effects are impaired by add-back of wild-type AMOT, but not of an AMOT mutant incompetent for YAP/TAZ binding (Fig. 5d and Extended Data Fig. 9e). Consistently, we also found that triple KO of AMOT in poorly aggressive, Ras-transformed human mammary epithelial cells (MII) potently enhanced their transformation in mechano-OFF conditions, as measured by mammosphere and soft-agar assays (Extended Data Fig. 9f,g) and confirmed by in vivo tumorigenesis assays (Fig. 5e). Coherently, we also demonstrated that the effects of AMOT depletion were strictly YAP/TAZ-dependent, as shown by concomitant AMOT and YAP/TAZ depletion or treatment with TEAD inhibitors (Extended Data Fig. 9f,g). In the opposite direction, we also proved that increasing the levels of wild-type AMOT, but not of AMOT incapable of YAP/TAZ association, impairs tumorigenic capacities of the highly aggressive MDA-MB-231 mammary cells. This was demonstrated in vitro, by inhibited soft-agar colony growth (Extended Data Fig. 9h), and in vivo, by impairment of tumor formation induced by experimentally supporting AMOT levels (Fig. 5f). The role of AMOT as candidate tumor suppressor in human tumors was then validated by digital pathology-based quantification of AMOT protein levels in a cohort of chemo-naïve TNBC samples vs. adjacent normal tissues. We found that AMOT is undetectable in TNBC, but present in the luminal compartment of the normal epithelium from which TNBC derive (Fig. 5g and Extended Data Fig. 10a,b). Importantly, in these patient samples, AMOT levels are invariably and quantitatively anti-correlated with YAP/TAZ nuclear localization (Fig. 5g and Extended Data Fig. 10a,b). We conclude from this collective set of biological assays that AMOT is a dominant determinant of YAP/TAZ activity, required and sufficient for cellular mechano-responses.

## Discussion

One of the most fascinating and least understood aspects of cell biology is how mechanical signals link the cell’s own shape and inner structure with gene expression and cell behavior. In recent years, several individual molecular steps - including the assembly of Focal adhesions ^8, 9^, actomyosin contraction ^5, 10^, nuclear deformation ^16, 17^, and YAP/TAZ entering through nuclear pores ^51^ – have been linked to mechanotransduction. Yet, this remains scattered knowledge as none of these discrete events can, by itself, explain mechanosignaling: it is the cell that integrates its response to forces across scales and organelles to determine its own mechanical state. Here we reveal that distinct subcellular structures are part of a mechanical continuum linking biophysical cues to YAP/TAZ mechanosignaling through control of AMOT stability by the centrosomal proteasome. We show that AMOT is a cardinal mechano-rheostat, serving as the long-sought cytoplasmic sink for the key mechanotransducing transcription factors YAP/TAZ.

We show that distinct cytoskeletal and cytoskeletal-associated structures act as an integrated whole to transduce forces. In particular, here we demonstrate the essential role of microtubules as pillars of cellular mechanosignaling, on par with, and downstream of, the F-actin cytoskeleton. MTs have been so far envisioned mainly as static determinants of cell shape, in virtue of their rigidity and ability to withstand compressive forces. In contrast, the direct contribution of MTs to mechanosignaling has remained poorly explored. Here we show that the cellular response to mechanical cues imposes a dramatic restructuring of the MTOC and of MT spatial organization (Fig. 6a,b). In mechanically inhibited cells, MTs are acentrosomal, with their minus end oriented at the cell periphery, and structured as a lattice that surrounds the nucleus. Microtubules radiating from the cell center in mechano-ON cells emanate from perinuclear centrosome generating a polarized astral arrangement that, in turn, facilitate further spreading from the Golgi complex ^63–66^, and from the perinuclear centrosome generating a polarized astral arrangement. The latter is essential for dynein-mediated transport of AMOT by direct binding of AMOT to the dynactin component DCTN1, at the level of its ICD.

**Fig. 6:**
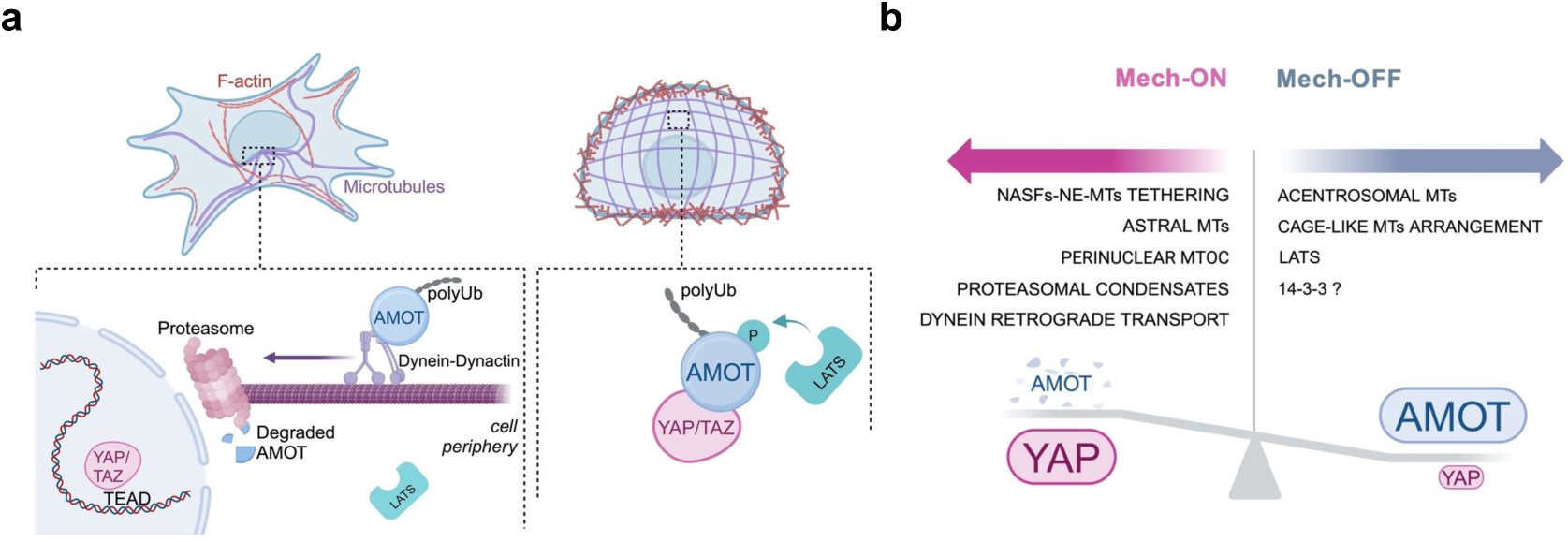
Schematic representation of the mechanical rheostat mechanism centered on the stability of AMOT proteins to control YAP/TAZ mechanoresponses. **a)** Top: schematics of the cytoskeletal organization of mechano-ON (left) vs. mechano-OFF (right) cells. Bottom: schematic representation of the mechano-signaling events controlling the stability of AMOT proteins. In mechano-ON cells AMOT protein is controlled by binding to the Dynein/Dynactin complex, allowing fast dynein-mediated transport of AMOT through the microtubular aster toward the pericentrosomal proteasome where AMOT is locally degraded. In mechano-OFF cells absence of a microtubular aster allows accrual of AMOT protein levels. Here, LATS kinase further contributes to AMOT stability by direct phosphorylation of AMOT, averting it from its degradation route. In mechano-OFF cells, increase of AMOT levels serves as cytoplasmic sink for YAP/TAZ, impeding nuclear accumulation. **b)** Schematic representation of factors and cytoskeletal components tipping the balance towards AMOT degradation and YAP/TAZ activation in mechano-ON cells are shown on the left and those that drive AMOT stabilization in mechano-OFF cells are shown on the right.

We propose that in mechano-ON cells, the centrosome, with its dual function of serving as MTOC and as physical scaffold for concentrating protein degradation, is a mechanoregulated cellular structure essential for YAP/TAZ regulation. We show that centrosome formation mediates the effects of rigidity sensing downstream of ECM adhesion and stress fiber formation. Three lines of evidence support this conclusion. First, specific disruption of radial MTs emanating from the centrosome in mechanically challenged cells prevents AMOT degradation and YAP/TAZ activation. Second, boosting stress fiber formation is inconsequential for YAP/TAZ activation in cells with inhibited radial MTs. Third, inducing MTOC formation in F-actin depleted cells by means of NLP1 overexpression is sufficient to restore AMOT degradation and YAP/TAZ nuclear entry. It is worth noting that, in mechano-OFF cells - the typical cell state of epithelial monolayers - increased F-actin cortical tension across cell-cell junctions may also influence microtubule orientation by repositioning MTOC components from the centrosome to apical or cortical structures, or by contributing to the stabilization of cortex-associated microtubule minus ends through γ-tubulin puncta ^67, 68^. These effects could at least facilitate the formation of acentrosomal microtubule arrays as cells experience decreasing levels of mechanical stimulation or arrange in monolayers and contribute to diminished AMOT delivery to the proteasome at the cell center. Curiously, we found that the two MT arrangements may in fact co-exist at intermediate levels of substrate stiffness, suggesting that these are alternative, but not mutually exclusive, modalities of MT organization.

The presently described AMOT regulation represents an energy-consuming but efficient system to readily shift a constantly produced YAP/TAZ inhibitor from degradation to accumulation. This warrants a swift adaptation of the cell to changes in its physical microenvironment. Since AMOT protein is constitutively tagged for degradation, we further note that its accrual in mechanically inhibited cells may occur by two mechanisms: accumulation of de novo synthesized AMOT at a rate exceeding its degradation, or by a yet unknown step in mechanosignaling, that is, AMOT deubiquitylation by the activity of specific deubiquitinating enzyme(s). Further work is required to distinguish between these possibilities.

An appealing aspect of our findings is to provide a unifying model that mechanistically merges the activity of the two main inputs feeding on YAP/TAZ activity, the Hippo cascade and mechanosignaling. The current model by which YAP/TAZ are regulated by Hippo kinases is through direct YAP/TAZ phosphorylation, whose inhibitory mechanisms remain incompletely defined, spanning from regulation of protein stability to tuning association with undefined cytoplasmic anchors ^48^. In the context of mechanosignaling, however, our data provide, at least in part, a departure from this model, as here we show that LATS phosphorylation of AMOT represents a dominant layer by which the Hippo cascade regulates YAP/TAZ. AMOT phosphorylation by LATS1/2 is constitutive: on the one hand, this is known to favor its association to 14-3-3 ^43^; on the other, as shown here, in mechano-ON cells this phosphorylation is skewing AMOT away from its cell shape-MTs-centrosome dependent degradation route. In other words, by hindering fast AMOT retrograde transport and degradation, LATS1/2 tip the balance to regenerate an AMOT pool. We speculate that AMOT salvage may be further reinforced by AMOT association to 14-3-3 ^43^. In other words, the effect of Hippo signaling on YAP/TAZ mechanotransduction appears overly indirect, by promoting AMOT stabilization and dampening the amplitude of mechanosensing. This clearly leaves open intriguing questions on how, and in which contexts, YAP/TAZ direct phosphorylation by LATS may further tune YAP/TAZ mechanoresponsiveness. This may occur in parallel to cell mechanics, as downstream of Hippo activation mediated by cell-cell junction, GPCR signaling and by polarity complexes in epithelial cells. These regulations may intersect with mechanosignaling shifting the thresholds of YAP/TAZ mechano- regulation.

Our data support the leading role of the nuclear envelope in cellular mechanotransduction ^16, 17, 22^. We show that the nuclear envelope serves as hub mediating the physical and tensional connectivity between the F-actin and MTs cytoskeletal systems through distinct LINC complexes. The role of NASF as preferred F-actin subpool to mechanically tether the nuclear envelope, here investigated primarily in epithelial cell types, may be further reinforced by more dorsal “actin-cap”-like fibers that are specific of other cell types, chiefly fibroblasts ^69^.

The central role of the nuclear envelope as cellular mechanical hub may also explain why damage to nuclear structural proteins results not only in decreased mechanical fitness of the whole cell ^16, 70^, but also in reduced mechanical activation of gene transcription ^17^. In this view, it is tempting to speculate that this may occur, at least in part, through AMOT stabilization and defective YAP/TAZ activation. Of note, laminopathies and LINC defects converge at inducing aging-related defects ^3^, and, intriguingly, the same is true for other ageing-like genetic disorders, such as mutations in Ninein (e.g. Seckel disease) ^71^, that impact on MTOC tethering to the nuclear envelope. Our data may thus offer a point of convergence between these apparently unrelated observations, particularly considering the recently discovered role of declining YAP/TAZ activity in ageing and senescence ^72^.

Recent studies have proposed that the nuclear entry of YAP/TAZ can be regulated by the permeability of nuclear pores ^51^. Clearly, the regulatory layer provided by the nuclear permeability model may co- exist with the molecular regulation of YAP/TAZ by cytoplasmic sinks, particularly through mechanisms centered on the stability of the AMOT protein as here proposed. We further note that the nuclear permeability model may contribute to YAP/TAZ regulation only in the mechano-ON cell state, as it can hardly explain the control of mechanosignaling in resting, mechano-OFF state. Our results instead point to an overarching regulation of YAP/TAZ cytoplasmic retention determined by AMOT levels. Indeed, in cells experiencing soft substrates - that is, cells with predicted less permissive nuclear pores - experimental loss of AMOT nevertheless massively increases YAP nuclear entry. Of note, the nuclear permeability model relates to a low- or zero-AMOT cell state, as it was proposed in studies in which cells were experiencing levels of extracellular matrix (ECM) rigidity (e.g., greater than 5 kPa) that we have shown are sufficient to deplete cells of their AMOT protein pool (Fig. 2c). That said, experimental stabilization of AMOT in mechano-ON cells is sufficient to effectively oppose YAP/TAZ nuclear entry despite more permissive nuclear pores. Taken together, these results paint a picture in which YAP/TAZ subcellular localization is dictated by AMOT stability in mechano-ON versus OFF states but can be further controlled by a second regulatory layer the level of nuclear pores in AMOT-low mechano-ON cells.

We speculate that AMOT availability in normal epithelia may spatially confine YAP/TAZ activation to specific locations and cell states, at the same time preventing unscheduled YAP/TAZ activation with its associated tumorigenic potential ^4^. Here we show that AMOT is a novel candidate tumor suppressor acting through YAP/TAZ inhibition, raising the tempting possibility that pharmacological treatments aimed at supporting AMOT stabilization may represent promising avenues of cancer therapy. Promise in this direction may come for example from probing the structural pockets here found to be required for AMOT-DCTN1 association. This may represent an orthogonal approach to current efforts aiming at blunting YAP/TAZ, so far entirely devoted to oppose YAP/TAZ at the nuclear level by TEAD-inhibiting drugs ^73^.

Finally, the presently described connections between centrosome formation, AMOT degradation and YAP/TAZ activation may also contribute to explain why numerical or structural centrosomal amplification is a hallmark of cancer and a marker of malignancy ^74^. Although this phenomenon has been historically connected to chromosome instability, the mechanisms by which centrosome amplification drives cancer remains elusive. Our data at least raise the possibility that this event may also boost cancer mechanosignaling and induce gain of YAP/TAZ-driven proliferation, invasiveness and plasticity. Thus, the centrosome may be an actionable target particularly for solid tumors whose malignancy is associated to increased stromal rigidity and YAP/TAZ activation.

## Acknowledgements

We thank colleagues sharing their plasmids on Addgene, Maria Vittoria Dieci, Valentina Guarneri, Matteo Fassan and Vincenza Guzzardo for human TNBC sample in the context of the METAMECH Consortium, and all members of our group for critical reading of the manuscript.

This research has received funding from the following agencies/charities: FONDAZIONE AIRC under 5 per Mille 2019 - ID. 22759 program, and under IG 2019 - ID. 23307 project to S.P.; the European Research Council Executive Agency (ERCEA) under the ERC-2022-ADG Grant Agreement n. 101098074-CHARTAGING to S.P.; FONDAZIONE AIRC under IG 2022 - ID. 27883 project to M.C., the European Union–NextGenerationEU and STARS@UNIPD, GF-MET-Growth factor signaling in metastatic organotropism to F.Z and the European Union - Next Generation EU, Mission 4, Component 2, CUP C93C22002780006, Spoke n.2 (“Cancer”) to F.Z., T.P and S.P. and Spoke n.5 (“Inflammatory diseases”) to M.C and G.B.

## Author Contributions

G.V. performed most of the experiments in vitro and in vivo, trained and received help from C.J.A., and contributed to the writing. A.C. and G.M. helped with the CRISPR KO clones and mice experiments.

A.G. and G.B. supported the mechanical bioassays. M.C. helped with data mining. M.D.P., E.P., L.S.P., A. S., and R.A.S. helped with protein structural studies. R.C. helped with 3D image reconstructions. T.P. developed the methodology for digital pathology quantitative analyses. F.Z. and P.C contributed with ideas and troubleshooting. S.P. and T.P. conceived the initial hypothesis and experimental design, organized the work and wrote the paper.

## Resource availability

All data are available in the manuscript or the supplementary materials. Correspondence and requests for materials should be addressed to Stefano Piccolo.

## METHODS

### Plasmids and reagents

LatrunculinA, CytochalasinD, Dynarrestin, Doxycycline, MG132 (Z-Leu-Leu-Leu-al), MG115 (Z-Leu-Leu-Norvalinal), Dasatinib and XAV939 were from Sigma. WIKI4 and JW55 were from EMD Millipore. Lactacystin was from Santa Cruz Biotechnology. Fasudil was from Tocris Bioscience. Defactinib, Vincristine, Nocodazole, Ciliobrevin D and Cycloheximide were from Selleckchem. VT107, GNE6640 and P005091 were from MedChemExpress. Rho inhibitor 1 (Exoenzyme C3 Transferase) was from Cytoskeleton, Inc. A complete list of plasmids employed is provided in Supplementary Table 4.

### Cell culture

MCF10A cells were a kind gift from F. Miller (Karmanos) and were cultured in DMEM/F12 (Gibco) with 5% horse serum, glutamine and antibiotics, freshly supplemented with insulin (Sigma-Aldrich), h- EGF (Peprotech), hydrocortisone (Sigma-Aldrich) and cholera toxin (Sigma-Aldrich). MCF10A-YAP-eGFP-KI cells were a kind gift from J.T.Liphardt ^75^, and were cultured as MCF10A. MII cells were a kind gift from S. Santner ^76^, and were cultured as MCF10A. HEK293 and HEK293T cells were from ATCC and were cultured in DMEM (Gibco) supplemented with 10% FBS, glutamine and antibiotics. U2OS cells were from ATCC and were cultured in DMEM (Gibco) supplemented with 10% FBS, glutamine and antibiotics. MDA-MB-231 cells were from the ICLC, and were cultured in DMEM/F12 (Gibco) supplemented with 10% FBS, glutamine and antibiotics. Bone marrow-derived hMSCs were purchased from Lonza and grown according to the manufacturer’s instructions. HEK293 LATS dKO cells were a kind gift from KL Guan ^77^, and cultured as HEK293.

NLP-expressing HEK293 cells were obtained by transduction with plenti-NLP-c-Myc-DDK-P2A-puro followed by puromycin selection, and were cultured in DMEM (Gibco) supplemented with 10% FBS, glutamine and antibiotics.

RFP-AMOT-expressing MCF10A-YAP-eGFP-KI cells were obtained by transduction with doxycycline-inducible lentiviral construct (pcw57.1-RFP-AMOTL1), and were cultured in DMEM/F12 (Gibco) with 5% horse serum, glutamine and antibiotics, freshly supplemented with insulin (Sigma-Aldrich), h-EGF (Peprotech), hydrocortisone (Sigma-Aldrich) and cholera toxin (Sigma-Aldrich). Doxycycline 2μg ml^-1^ was administered to cells 24 hours before the experiments to induce RFP-AMOT overexpression.

HEK293 AMOT tKO cells were obtained by CRISPR-Cas-mediated gene editing, by transfection of pSpCas9(BB)-2A-Puro-AMOTtKO-A or pSpCas9(BB)-2A-Puro-sgGFP as negative control, using TRANS-IT LT1 transfection reagent following manufacturer instruction. Cells were selected by puromycin and after 3 weeks independent clonal cell lines (control, AMOT tKO#1 and AMOT tKO#2) were picked, expanded and tested by immunoblotting. HEK293 AMOT tKO cells were cultured as HEK293 cells.

HA-AMOT^wt^ and HA-AMOT^L/PPXA^ -expressing MDA-MB-231 cells were obtained by transduction with doxycycline-inducible lentiviral constructs (pcw57.1-HA-AMOT^wt^ or pcw57.1-HA-AMOT^L/PPXA^), and were cultured in DMEM/F12 (Gibco) supplemented with 10% Tetracycline-free FBS, glutamine and antibiotics. Doxycycline 2μg ml^-1^ was administered to cells 24-48 hours before the experiments to induce HA-AMOT^wt^ or HA-AMOT^L/PPXA^ expression.

MII AMOT tKO cells were obtained by CRISPR-Cas-mediated gene editing by nucleofection using Amaxa 4D nucleofector (Lonza) with pSpCas9(BB)-2A-Puro-AMOTtKO-B or pSpCas9(BB)-2A-Puro- sgGFP as negative control. Cells were selected by puromycin and after 3 weeks independent clonal cell lines (control, AMOT tKO#1 and AMOT tKO#2) were picked, expanded and tested by immunoblotting. MII AMOT tKO cells were cultured as MII cells.

AMOT-expressing hMSCs were obtained by transduction with doxycycline-inducible lentiviral constructs pcw57.1-HA-AMOT^wt^ and were treated with Doxycycline 2μg ml^-1^ throughout all the duration of adipogenic differentiation assays.

RFP-AMOT expressing HEK293 cells were obtained by transduction with doxycycline-inducible lentiviral constructs (pcw57.1-RFP-AMOTL1), and were cultured in DMEM (Gibco) supplemented with 10% Tetracycline-free FBS, glutamine and antibiotics. Doxycycline 0,33 μg ml^-1^ was administered to cells 24 hours before the experiments to induce mild RFP-AMOT expression.

FLAG-YAPwt and FLAG-YAP5SA expressing MCF10A cells were obtained by transduction with doxycycline-inducible lentiviral constructs (pcw57.1-FLAG-YAP^wt^ or pcw57.1-FLAG-YAP^5SA^), and were cultured as MCF10A cells. Doxycycline 0,04 and 0,2μg ml^-1^ was administered to cells 24-48 hours before the experiments to induce comparable expression of FLAG-YAPwt and FLAG-YAP5SA, respectively.

All parental cell lines were authenticated by BMR Genomics. All cell lines were routinely tested to exclude mycoplasma contamination.

#### RNA interference and DNA transfections

siRNA transfections were performed with Lipofectamine RNAi-MAX (Thermo Fisher Scientific) in antibiotic-free medium according to the manufacturer’s instructions. Sequences of siRNAs are provided in Supplementary Table 5. Validations of siRNAs efficiency are provided in Supplementary Figure 2. DNA transfections were performed with TransitLT1 (Mirus Bio) according to the manufacturer’s instructions. For dual siRNA/DNA transfections, DNA transfections were performed 8 hours after siRNA transfection. For dual siRNA transfections, transfections were performed 8 hours apart. Cells were harvested 48 hours post siRNA or DNA transfection, unless differently specified.

#### Human mammary tissue

Discard tissue from anonymized healthy women undergoing reduction mastoplasty surgery and human TNBC archival samples were collected with informed consent according to our institutional guidelines and the Azienda Ospedaliera di Padova Ethics Committee (CESC).

#### Human mammary luminal cell isolation and culturing

Single-cell suspensions of primary human mammary cells were generated as previously described ^49^, with minor modifications. Briefly, the ductal tree was mechanically minced and enzymatically digested in tissue dissociation medium (Advanced DMEM-F12 supplemented with HEPES, 1.5% GlutaMAX, 600 U ml^−1^ collagenase and 200 U ml^−1^ hyaluronidase at 37 °C overnight). Cells were spun down 3 min at 700 r.p.m., and the pellet was further dissociated in 0.25% trypsin-EDTA for 5 min followed by the addition of 5 μg ml^−1^ dispase and 1 μg ml^−1^ DNaseI for a further 10 min. Digestion was stopped in Advanced DMEM 10% FBS, and cells were filtered through a 40 μm strainer to remove residual tissue fragments and cell aggregates. Single-cell suspensions of primary mammary cells were stained with CD31 (BioLegend no. 303119), CD45 (BD Biosciences no. 557833), CD49f (BD Biosciences no. 555736) and EpCAM (BD Biosciences no. 347197) in DMEM for 30 min at 4 °C. After excluding CD31+CD45+(Lin+) cells, mammary cells were sorted into four populations, as previously shown ^49^: Luminal differentiated cells, CD49f–EpCAM+; Luminal progenitor cells, CD49f+EpCAM+; basal cells, CD49f+EpCAM−; and stromal cells, CD49f−EpCAM−, using a FACS Aria III (BD Biosciences). Freshly isolated primary Luminal differentiated cells were seeded in human mammary epithelial growth medium (HMGM; Advanced DMEM/F12 supplemented with HEPES, GlutaMAX, 0.5% FBS, 4 μl ml^−1^ BPE, 10 ng ml^−1^ hEGF, 10 μM Y27632, 10 μM forskolin and antibiotics). After FACS purification, cells were seeded in 24-well plates coated with collagen I and transduced with lentiviral vectors (virus suspension was mixed 1:1 with medium). For primary mammary cell reprogramming, cells were transduced for 48 hours with FU-tetO-HER2-CA in combination with rtTA-encoding lentiviruses (FudeltaGW-rtTA) and with FU-tetO-AMOT^wt^ or FU-tetO-AMOT^L/PPXA^. As a (negative) control, cells were transduced with EGFP-expressing vector (FUW-tetO-EGFP) in combination with rtTA-encoding lentiviruses. After infection, adherent cells were washed and treated with 2 μg ml^−1^ doxycycline for 7 days in HMGM to activate tetracycline-inducible gene expression. After 1week, mammary cells were detached with trypsin and seeded at a density of 2,000 cells per well in 24-well ultralow attachment plates (Corning) in mammary clonogenic suspension medium (Advanced DMEM/F12 containing HEPES, GlutaMAX, antibiotics, 5% Matrigel, 2% FBS, 10 ng ml^−1^ human EGF, 4 μl ml^−1^ BPE, 10 μM forskolin and 2 μg ml^−1^ doxycycline). Primary colonies were counted 14 days after seeding.

#### Orthotopic mammary fat pad transplantations

MII, MII-AMOTtKO, MDA-MB-231 and AMOT-expressing MDA-MB-231 cellular suspensions (1x10^6^ / mice) in 50% Matrigel / PBS1x were injected into the inguinal mammary fat pads of female NOD-SCID mice (IMSR no. CRL:394, Charles River) at 3 weeks of age. For experiments of Fig. 5f, animals were then administered doxycycline (2 mg ml^-1^ in water supplemented with 10 mg ml^-1^ sucrose) in their drinking water for 4 weeks. 8 weeks (for MII cells) and 4 weeks (for MDA-MB-231 cells) after injection, mice were sacrificed to extract tumor masses, and tumor mass volume was measured by caliper measurement of the three major axes. Animal experiments were performed adhering to our institutional guidelines as approved by the animal welfare body (Organismo Preposto al Benessere Animale; OPBA).

#### Microfabrications

Fibronectin-coated hydrogels were obtained as previously described ^78^ with minor modifications. Hydrogel formulations in AA (Acrylamide), BA (bis-acrilamide), TEMED (tetramethylethylenediamine) and APS (ammonium persulfate) were as follows: 3.5% w./v. AA, 0.03% w./v. BA, 1:1,000 v./v. TEMED and 0.1% w./v. APS (0.7 kPa); 3% w./v. AA, 0.15% w./v. BA, 1:1,000 v./v. TEMED and 0.1% w./v. APS (3 kPa); 5% w./v. AA, 0.225% w./v. BA, 1:1,000 v./v. TEMED and 0.1% w./v. APS (13 kPa); and 8% w./v. AA, 0.48% w./v. BA, 1:1,000 v./v. TEMED and 0.2% w./v. APS (50 kPa). In all the compositions, AA% represents the sum of acrylamide and N-hydroxyethyl acrylamide, which is fixed at 0.1M. Cells (4 × 10^4^ cells per cm^2^) were seeded in a drop of complete culture medium on top of fibronectin-coated hydrogels; after attachment, the hydrogel-containing wells were filled with the appropriate culture medium. Cells were collected for immunofluorescence, RNA, protein extraction or Luciferase assays after 24h.

Micropatterned glass substrates were prepared by ultraviolet (UV) photolithography as previously described ^79^, with minor modifications. Briefly, a positive photoresist (S1805 G2, Microchem) was spin coated on glass coverslips, previously silanized with 3-(trimethoxysilyl) propyl methacrylate (Sigma-Aldrich). The film was then patterned by UV exposure (collimated lamp at 365 nm) through a brightfield chrome photomask and developed with MF-319 (Microchem), achieving squares of photoresist. Linear polyacrylamide was then grafted on the patterned surface of the glass, polymerizing an 8% w/v solution of acrylamide in milliQ water with 0.0225% w/v of ammonium persulfate and 15:1000 v/v of TEMED. The substrates were left overnight in water to wash out unreacted reagents. Finally, the resist was stripped by dipping the substrates in DMSO and milliQ water; the coverslips were UV sterilized and the adhesive islands were functionalized incubating a 10 μg ml^−1^ fibronectin solution for 1 h at 37 °C. Micropatterned coverslips were washed twice in PBS and 1x10^5^ cells (MCF10A, U2OS, HEK293) per micropattern were seeded in complete culture medium. Cells were gently washed twice with complete medium 2 hours after seeding to remove unattached cells. Cells were fixed after 24 hours for immunofluorescence.

Microfabricated fibronectin islands employed in experiments of Fig. 5b and Extended Data Fig. 9d were purchased from Cytoo SA. MCF10A (1x10^6^) cells were plated in a 35mm dish containing a single Cytoo glass slide, and nonadherent cells were washed with medium after 2 hrs. After 24 hours, cells were processed for proliferation assay, as indicated by the manufacturer (BrdU Cell proliferation kit Merck). Briefly, cells were incubated for 1 hour with a pulse of BrdU to label cells undergoing DNA duplication. Cells were fixed and processed for anti-BrdU immunofluorescence (αBRDU). Pictures of BrdU stainings of 50 individual multicellular islands were turned into binary signals, superimposed to obtain the relative pixel frequency, and color-coded using ImageJ (BrdU stack). Pixel values indicate the percentage of cells in individual pictures displaying BrdU positive signal at that location. The same process was applied to pictures of nuclei (stained with Hoechst), ensuring a uniform distribution of cells within the monolayers. Proliferation rates in each sample were mapped with Fiji-ImageJ by plotting BrdU pixel frequencies along the X-axis of an imaging window set at the center of the Y-axis. Confocal images were obtained with a Leica SP5 microscope and analyzed using Fiji-ImageJ.

#### Immunofluorescence

Immunofluorescence on PFA-fixed cells was performed as previously described ^62^. Primary antibodies were YAP (Santa Cruz Biotechnology, sc-101199) previously validated as YAP/TAZ antibody ^5, 10, 49, 80^, LaminA/C (Santa Cruz Biotechnology, sc-376248), Paxillin (Abcam no. ab32084), GFP (Abcam ab13970), Proteasome subunit alpha 5 (PSMA5) (Origene, TA332887), AMOT (Santa Cruz Biotechnology, sc-166924), AMOTL1 (Sigma HPA001196), alpha Tubulin (Abcam, ab18251), γ- Tubulin (Santa Cruz Biotechnology, sc-17787), acetylated-Tubulin (ab24610), Pericentrin 1 (Santa Cruz Biotechnology, sc-376111), LaminC/C (Abcam, ab8984), RNF146 (Thermofisher, H00081847-B01P). F-actin was stained with Alexa Fluor 568 Phalloidin (Invitrogen). Secondary antibodies (1:200) were from Invitrogen. Samples were counterstained with Hoechst to label cell nuclei. Slides were mounted using Prolong Diamond (Sigma).

For proteasomal staining (Fig. 2a,b) cells were treated with Lactacystin 10 μM 6 hours before harvesting. For microtubules and centrosomal staining, cells were fixed as previously described ^81^ with minor modifications: cells were fixed and permeabilized in a solution of 3% PFA, 0,2% glutaraldehyde and 0,25% Triton-X 100 in PBS 1x for 15 minutes at room temperature, and then washed twice in PBS 1x for 10 minutes. Before staining, cells were treated with a quenching solution (10mM MES, 150 mM NaCl, 5 mM EGTA, 5 mM MgCl_2_ and 5 mM glucose, pH 6.1) freshly added with 1mg ml^-1^ sodium borohydride for 15 minutes on ice, and then washed with PBS 1x. For a conformation-sensitive staining of the nuclear lamina (LmnA C-C staining), cells were fixed in 2% PFA in PBS 1x. Unless otherwise specified, confocal images were obtained with a Leica Stellaris 5 microscope and analyzed using LASX software and Fiji-ImageJ. Super-resolution images of cytoskeletal staining (Fig. 1e,g,k,l; Fig. 3j and Extended Data Fig. 2b; Extended Data Fig. 3a,d and Extended Data Fig. 4c) were obtained with the Leica Stellaris 5 LIGHTNING module. YAP/TAZ nuclear versus cytoplasmic ratios was quantified by using Fiji-ImageJ analysis software with the same threshold in each staining. γTURC numbers were quantified as the number of γTubulin positive dots in each cell by using Fiji-ImageJ analysis software. Proteasome puncta were quantified as the number of 20S positive spots in each cell by using Fiji-ImageJ analysis software. FAs were measured as the average length of the major axis of paxillin-positive signal per cell by using Fiji-ImageJ analysis software. The apical-to-basal lamin intensity ratios (Extended Data Fig. 3b,c) were determined as previously described ^23^ using Fiji-ImageJ analysis software. 3D reconstructions of confocal images were obtained using Z-stacks with LASX software (Leica). The 3D reconstructions shown in Fig. 1i,j and Supplementary Video 2 were obtained with AIVIA (Leica) software, by training and applying a machine-learning pixel classifier algorithm to differentially segment phalloidin-stained F-actin fibers that directly contact the nuclear lamina (NASFs, shown in orange in Fig. 1i,j and Supplementary Video 2) and those that do not (shown in grey in Fig. 1i,j and Supplementary Video 2).

#### Immunohistochemistry

Immunohistochemical staining experiments were performed on PFA-fixed, paraffin wax-embedded tissue sections as previously described ^61^. For immunohistochemistry the antibodies used were anti-YAP (CST #4912), anti-TAZ (Sigma-Aldrich HPA007415) and anti-AMOTL1 (Sigma-Aldrich HPA001196). Brightfield images were obtained with a Nanozoomer Scanner 2.0RS (Hamamatsu), equipped with the NDPscan3.1 software. Immunohistochemical stainings were quantified with QuPath 5.0 ^82^, by adapting a StarDist ^83^ deep-learning-based cell segmentation algorithm. Relative AMOT protein abundance and YAP/TAZ nuclear-to-cytoplasmic ratios were normalized and expressed as Z-scores for comparative analysis.

#### Live imaging

For live imaging experiments of Fig. 2f, RFP-AMOT expressing MCF10A-YAP-eGFP-KI cells were seeded on μ-dishes (Ibidi) previously coated with fibronectin 20 μg ml^-1^ (Santa Cruz Biotechnology) in 0,33μg ml^-1^ doxycycline-containing medium 24 hours before live imaging.

For live imaging experiments of Supplementary Video 3, RFP-AMOT expressing HEK293 cells were seeded on μ-dishes (Ibidi) previously coated with fibronectin 20 μg ml^-1^ (Santa Cruz Biotechnology) in 0,33μg ml^-1^ doxycycline-containing medium 24 hours before live imaging. The day after seeding, cells were kept in complete medium supplemented with Dynarrestin 10μM or with DMSO as negative control starting from 3 hours before live imaging. For microtubules visualization, Tubulin-555 (SpyroChrome) wasSpy diluted in culture medium according to manufacturer’s instructions 1 hour before live imaging. All live imaging was acquired with a Leica Stellaris5 confocal microscope equipped with an Okolab incubator chamber. 3D time lapse videos were reconstructed using AIVIA software (Leica) by training and applying a machine-learning pixel classifier algorithm to differentially segment AMOT aggregates and microtubules.

#### In situ Proximity Ligation Assay (PLA)

In situ PLAs were performed with Duolink in situ reagents (Sigma-Aldrich) on PFA fixed samples, according to manufacturer’s instructions. Antibodies used were YAP1 (Proteintech, 13584-AP) and AMOT (Santa Cruz Biotechnology, sc-166924). Samples were counterstained with Hoechst to label cell nuclei. Slides were mounted using Prolong Diamond (Sigma). Images were acquired with a Leica Stellaris5 confocal microscope and analyzed using Fiji-ImageJ software.

#### hMSC adipogenic assay

Adipogenic differentiation was performed as previously described ^5^, with minor modifications. For the experiments shown in Fig. 5a and Extended Data Fig. 9a,b , hMSCs were transfected with the indicated siRNAs and reseeded on micropatterns after 24 hours; cells were then cultured for 5 days in adipogenic induction medium (Lonza) followed by 1 day in adipogenic maintenance medium (Lonza) according to manufacturer instructions, until harvesting at day 6 of differentiation. Where indicated, cells were treated with Cyto.D 500 nM throughout the experiment. For the experiments shown in Extended Data Fig. 9c, hMSCs were transduced with pCW57.1-MCS (Empty vector) or pCW57.1-HA-AMOTwt lentiviral constructs and then cultured for 5 days in adipogenic induction medium (Lonza) followed by 1 day in adipogenic maintenance medium (Lonza) according to manufacturer instructions in presence of Doxycycline 2 μg ml^-1^, until harvesting at day 6 of differentiation. Adipogenic differentiation was assayed by Oil Red staining (Sigma) and quantified as the Oil Red-positive area normalized to the number of cells (Hoechst-positive nuclei) using Fiji-ImageJ analysis software. Cells were treated as above and harvested after 24 hours of culture in adipogenic induction medium to measure YAP/TAZ activity (Extended Data Fig. 5n).

#### qRT-PCR

Cells were collected using the RNeasy Mini Kit (Qiagen) for total RNA extraction, and contaminant DNA was removed by DNase treatment (Qiagen). qRT-PCR analyses were carried out on reverse-transcribed cDNAs with QuantStudio 5 (Applied Biosystems, Thermo Fisher Scientific) and analyzed with QuantStudio Design and Analysis software (version 1.4.3). Expression levels are always normalized to GAPDH. PCR oligonucleotide sequences are listed in Supplementary Table 6.

#### Western blots

Immunoblots were carried out as previously described ^61^. Primary antibodies used are listed in Supplementary Table 7. Quantifications of relative protein levels from immunoblot bands were performed with Fiji-ImageJ analysis software, relative to the corresponding GAPDH band from the same lane. Uncropped immunoblot images are provided in Supplementary Figure 3.

#### Co-immunoprecipitation of endogenous proteins

Pulldown of endogenous AMOT (Fig. 3c,f), of endogenous DCTN1/p150-Glued (Fig. 3i, Fig. 4c and Extended Data Fig. 7n), and of endogenous YAP (Extended Data Fig. 7h) were performed on HEK293T cell lysates. For DCTN1/p150-Glued pulldown, cells were transiently transfected with 10ng cm^-2^ pCDNA3-HA-hAMOTwt or pCDNA3-HA-hAMOT-L178W for experiments in Fig. 3i and Extended Data Fig. 7n, and with 0,2ng cm^-2^ pCDNA3-HA-hAMOTwt or pCDNA3-HA-hAMOT-S175A for experiments in Fig. 4c and Extended Data Fig. 8d. Cells were re-plated at low confluency 24 hours post DNA transfection. Cells were treated with Dynarrestin 10μM for 6 hours before harvesting for experiments shown in Fig. 4c and Extended Data Fig. 7n and Extended Data Fig. 8d, and with proteasome inhibitor Lactacystin 10μM for 6 hours before harvesting for experiments shown in Fig. 3f. All samples were harvested in Marais buffer (20mM HEPES pH 7.8, 400mM KCl, 5mM EDTA, 0,4% NP-40, 10% Glycerol) freshly supplemented with 1mM DTT, protease (Sigma) and phosphatase (Roche) inhibitors and kept on ice. Samples were sonicated and cleared by centrifugation for 10 minutes at full speed at +4°C. Samples were diluted 1:7 v/v (for AMOT pulldown) or 1:3 v/v (for DCTN1/p150-Glued pulldown) in correction buffer (20mM HEPES pH 7.8, 3,1mM MgCl_2_, 3,5% Glycerol) freshly supplemented with protease and phosphatase inhibitors. For YAP pulldown, samples were harvested in YAP lysis buffer (20mM HEPES ph 7.6, 50 mM KCl, 0.1% NP-40, 0.1% Triton-X100, 5% glycerol, 5 mM MgCl_2_,1mM DTT, 1mM ATP, 1mM creatin-phosphate) freshly supplemented with protease and phosphatase inhibitors. Samples were mechanically disrupted by repeated passages through a 26G needle and cleared by centrifugation for 10 minutes at full speed at room temperature. Samples were then added of 100 mM NaCl, 1.5 mM MgCl_2_, 0.2 mM EDTA, 15% glycerol. Protein-G or protein-A coated magnetic Dynabeads (Thermofisher) were coated with 2μg of anti-AMOT (Santa Cruz Biotechnology) or anti- DCTN1/p150-Glued (Abcam) or anti YAP1 (Proteintech) antibody per sample and cell lysates were incubated with coated magnetic beads for 3 hours at 4°C (AMOT and DCTN1 pulldowns) or for 3 hours at room temperature (YAP pulldown) in gentle rotation. Beads coated with normal mouse IgG (Santa Cruz Biotechnologies) or with normal rabbit IgG (Sigma) were used as negative control for AMOT or DCTN1/p150-Glued and YAP pulldown, respectively.

#### Co-immunoprecipitation of overexpressed proteins

Pulldown of FLAG-DCTN1wt and FLAG-DCTN1-ICD-4Wmut shown in Fig. 3h and Extended Data Fig. 7m, pulldown of HA-hAMOT-S175E and HA-hAMOT-S175A shown in Fig. 4b and Extended Data Fig. 8c were performed on HEK293T cell lysates. For FLAG-DCTN1 pulldown, cells were transiently transfected with 60ng cm^-2^ pCDNA3-HA-hAMOTwt and 100ng cm^-2^ pCDNA3-FLAG-DCTN1wt or pCDNA3-FLAG-DCTN1-ICD-4Wmut. For HA-AMOT pulldown, cells were transiently transfected with 25ng cm^-2^ pCDNA3-HA-hAMOT-S175A or 12,5ng cm^-2^ pCDNA3-HA-hAMOT-S175E.Cells were re-plated at low confluency 24 hours post DNA transfection. The next day, cells were treated with Dynarrestin 10μM (for FLAG-DCTN1 pulldown) or with Lactacystin 10μM (for HA-AMOT pulldown) for 6 hours before harvesting. All cells were harvested in Marais buffer (20mM HEPES pH 7.8, 400mM KCl, 5mM EDTA, 0,4% NP-40, 10% Glycerol) freshly supplemented with 1mM DTT, protease and phosphatase inhibitors and kept on ice. Samples were sonicated and cleared by centrifugation for 10 minutes at full speed at +4°C. Samples were diluted 1:3 v/v (for FLAG-DCTN1 pulldown) or 1:7 v/v (for HA-AMOT pulldown) with correction buffer (20mM HEPES pH 7.8, 3,1mM MgCl_2_, 3,5% Glycerol), freshly supplemented with protease and phosphatase inhibitors.

Protein-G or protein-A coated magnetic Dynabeads (Thermofisher) were coated with 2μg anti-FLAG M2 (Sigma) or anti-HA (Abcam) antibody per sample and cell lysates were incubated with coated magnetic beads for 3 hours at 4°C in gentle rotation. Beads coated with normal mouse IgG (Santa Cruz Biotechnologies) or with normal rabbit IgG (Sigma) were used as negative control for FLAG or HA pulldown, respectively.

#### PARylation assays

Immunoprecipitation of FLAGmAMOTL1 shown in Extended Data Fig. 6e was performed on HEK293T cell lysates. Cells were transiently transfected with 1,5 ng cm^-2^ pCMV6-FLAG-mAMOTL1. 24hours post DNA transfection, cells were re-plated in sparse (5x 10^4^ cells per cm^2^) versus dense (2,5 x 10^5^ cells per cm^2^) conditions with proteasome inhibitors MG115 and MG132 (10μM). After 24 hours, cells were lysed in PAR buffer (50mM Tris-HCl pH 7,4; 150mM NaCl; 25mM sodium pyrophosphate; 1% NP40; 5% glycerol) freshly supplemented with 1μM ADP-HPD (Millipore), 1mM DTT, protease and phosphatase inhibitors and kept on ice. Samples were sonicated and cleared by centrifugation for 10 minutes at full speed at +4°C. Protein-G coated magnetic Dynabeads (Thermofisher) were coated with 2μg anti-FLAG M2 antibody (Sigma) per sample in PAR buffer for 1hour at room temperature and then used for FLAG-mAMOTL1 pulldown. Cell lysates were incubated with coated magnetic beads for 4 hours at 4°C in gentle rotation.

#### Ubiquitylation assays

Immunoprecipitation of FLAGmAMOTL1 shown in Extended Data Fig. 6f was performed on HEK293T cell lysates. Cells were transiently transfected siRNAs and after 24 hours with 1,5 ng cm^-2^ pCMV6- FLAG-mAMOTL1 and with 60ng cm^-2^ pRK5-HA-Ubiquitin-WT. Cells were re-plated 8 hours post DNA transfection in sparse (5x 10^4^ cells per cm^2^) or dense (2,5 x 10^5^ cells per cm^2^) conditions. After 15 hours, cells were treated with Lactacystin (10μM) for 6 hours and then were lysed in Ub-buffer (50mM HEPES pH 7,8; 200mM NaCl; 5mM EDTA; 1% NP40; 5% glycerol) freshly supplemented with freshly complemented with 250 ng/ml ubiquitin aldehyde (Millipore, 662056), 1mM DTT, protease and phosphatase inhibitors, and kept on ice. Samples were sonicated and cleared by centrifugation for 10 minutes at full speed at +4°C. Protein-G coated magnetic Dynabeads (Thermofisher) were coated with 2μg anti-FLAG M2 antibody (Sigma) per sample in Ub-buffer for 1hour at room temperature and then used for FLAG-mAMOTL1 pulldown. Cell lysates were incubated with coated magnetic beads for 4 hours at 4°C in gentle rotation in Ub-buffer supplemented with 2mM MgCl_2_. Stringent washes were then performed using Ub-wash buffer (50mM HEPES pH 7,8; 500mM NaCl; 5mM EDTA; 1% NP40; 5% glycerol) freshly complemented with 250ng/ml ubiquitin aldehyde, 1mM DTT, protease and phosphatase inhibitors.

#### Filamentous actin quantitation assays

To determine the amount of filamentous actin (F-actin) versus free globular-actin (G-actin) content in mechano-ON versus mechano-OFF cells (Extended Data Fig. 5i), MCF10A cells were seeded on stiff (40kPa) versus soft (0.7 kPa) hydrogels (2x10^^6^ total cells for each condition). After 24 hours cells were harvested and processed for quantitation of F-actin and G-actin cellular fractions by Western blot, using the G-Actin/F-actin In Vivo Assay Biochem Kit (Cytoskeleton, Inc. #BK037), according to manufacturer’s instructions.

#### Luciferase reporter assays

Luciferase assays were performed as previously described ^49^, with minor modifications. Briefly, the TEAD luciferase reporter 8xGTIIC–Lux (75 ng cm^−2^) was transfected together with phRG-TK-Renilla (25 ng cm^−2^) to normalize for transfection efficiency. Cells plated at low confluency were transfected with DNA or with both siRNAs and DNA, and replated in Mech.ON versus Mech.OFF conditions, as indicated. For experiments shown in Fig. 4d and Extended Data Fig. 5o, cells were re-plated at low confluency in Cyto.D 200nM containing medium. Cells were harvested 24 hours post re-plating. Firefly luciferase activity was measured with an Infinite F200PRO plate reader (Tecan). Data are presented as firefly/*Renilla* luciferase activity.

#### Mammosphere assays

MII control and AMOT tKO cells were seeded (2x10^3^ cells per cm^2^) as single-cell suspension in ultra-low attachment plates (Corning). Cells were cultured in DMEM/F12 supplemented with 0,4% BSA, antibiotics, 5 μg ml^-1^ insulin (Sigma-Aldrich), 10 ng ml^-1^ hEGF (Peprotech), 20 ng ml^-1^ bFGF (Peprotech) and 4 μg ml^-1^ Heparin. Where indicated, siRNA transfection was performed the day before mammosphere seeding. For the experiments in Extended Data Fig. 9g, cells were treated with DMSO or VT107 6μM 48h before mammosphere seeding, and treatments were refreshed 3 μM every 48 hours. Mammospheres were counted after 15 days.

#### Agar colony formation assays

MII cells were transfected with the indicated siRNAs and embedded in agar after 24 h. Effective YAP/TAZ downregulation was assessed by immunoblotting on the same cell population. Where indicated, MII cells were treated with DMSO or the TEAD inhibitor VT107 6 μM 48h before agar embedding. For the experiments in Extended Data Fig. 9h, MDA-MB-231 cells were treated with Doxycycline 2 μg ml^-1^ 24 hours before agar embedding. Clonogenic assays were performed as previously described ^80^; briefly, cells (1x10^4^) were resuspended in complete growth medium with 0.3% agarose (BD Biosciences) and were layered onto 0.6% agar beds in six-well plates. Complete medium was added on top of the cells and was replaced with fresh medium twice a week for 4 weeks. For the experiments in Extended Data Fig. 9g, DMSO or the TEAD inhibitor VT107 3 μM were added to fresh medium. For the experiments in Extended Data Fig. 9h, MDA-MB-231 cells were maintained in Doxycycline 2 μg ml^-1^ supplemented complete medium.

#### Lentiviral preparations

Lentiviral particles were prepared as previously described ^61^. Briefly, HEK293T cells were transiently transfected with lentiviral vectors (167 ng cm^-2^) together with packaging vectors pMD2-VSVG (42 ng cm^-2^) and pPAX2 (125 ng cm^-2^) using TransIT-LT1 (Mirus Bio) according to the manufacturer’s instructions.

#### Statistics

The number of biological and technical replicates and the number of animals is indicated in the Fig. Legends, Main text and Methods section. All tested animals were included. The animal ages and sexes are specified in the text and Methods section. The sample size was not predetermined. Randomization was not applicable to our experiments with cell lines. The Student’s *t*-test and ANOVA analyses were performed, as indicated in the captions of the figures and supplementary figures, with GraphPad Prism 8.0.2 for Mac software.

## SUPPLEMENTARY MATERIAL

- Supplementary Tables 1-7.

- Supplementary Videos 1-3.

- Supplementary Figures 1-3.

## Supplementary Tables Legends

**Supplementary Table 1.** List of common YAP/TAZ interactors, as retrieved from published protein-protein interaction datasets (BioGRID 4.4) and summary of immunoblot experiments testing the protein levels of each YAP/TAZ cytoplasmic interactor in cells subjected to mechano-ON versus mechano-OFF conditions.

**Supplementary Table 2.** List of AMOT proximal proteins and associated significant GO terms (p-value(-Log10) ≥ 2.6), as in C. D. Go et al.^38^.

**Supplementary Table 3.** List of common AMOT/AMOTL1/AMOTL2 interactors, as retrieved from published protein-protein interaction datasets (BioGRID 4.4).

**Supplementary Table 4.** List of plasmids employed in this study.

**Supplementary Table 5.** List of siRNA and gRNA sequences employed in this study.

**Supplementary Table 6.** List of qRT-PCR oligo sequences employed in this study.

**Supplementary Table 7.** List of primary antibodies employed in Western Blot analysis.

**Supplementary Table 8.** Numerical source data giving rise to graphical representations in the paper.

## Supplementary Videos Legends

**Supplementary Video 1. Microtubule architecture in mechano-ON versus mechano-OFF cells.**

Representative all-around view of the same 3D immunofluorescence reconstruction of MCF10A cells seeded in mechano-ON (40 kPa hydrogel) conditions, or in different mechano-OFF conditions (0.7 kPa hydrogel; Small, 100 μm^2^ micropatterns; dense, 2,5 x 10^5^ cells per cm^2^), as in Extended Data Fig. 1a. MTs were labelled by α-Tubulin staining (αTub, magenta) and nuclei were counterstained with Hoechst (cyan).

**Supplementary Video 2. Reconstruction of F-actin and nuclear envelope in mechano-ON versus mechano-OFF cells.**

Representative wrap-around view of the same 3D immunofluorescence reconstruction of MCF10A cells seeded in mechano-ON (40 kPa hydrogel), or in mechano-OFF (0.7 kPa hydrogel) conditions, as in Fig. 1i. The nuclear lamina was labelled by LaminA staining (LmnA, cyan), F-actin was labelled by Phalloidin staining (F-actin, grey). Nuclear associated stress fibers (NASFs) are highlighted in orange in the 3D reconstruction (see methods).

**Supplementary Video 3. Time lapse imaging of AMOT retrograde transport along MTs** Representative time lapse 3D reconstruction showing AMOT retrograde transport (right to left) along microtubules in cells seeded in mechano-ON conditions, either left untreated or treated with with Dynarrestin 10μM for 6hrs. MTs were labelled by SPY-Tubulin (purple) and RFP-AMOT was labelled in yellow (see methods).

## Supplementary Figures Legends

**Supplementary Figure 1: Quantifications of AMOT immunoblots shown in Fig. 2 and Extended Data Fig. 5**.

**a)** Quantification of AMOT protein levels in the immunoblot shown in Fig. 2c.
**b)** Quantification of AMOT protein levels in the immunoblot shown in Fig. 2d.
**c)** Quantification of AMOT protein levels in the immunoblot shown in Extended Data Fig. 5d.
**d)** Quantification of AMOT protein levels in the immunoblot shown in Extended Data Fig. 5f.
**e)** Quantification of AMOT protein levels in the immunoblot shown in Extended Data Fig. 5g.
**f)** Quantification of AMOT protein levels in the immunoblot shown in Extended Data Fig. 5h.
**g)** Quantification of AMOT protein levels in the immunoblot shown in Extended Data Fig. 5j.

**Supplementary Figure 2: Validations of siRNAs specificity.**

**a)** Quantitative real-time PCR (qRT-PCR) assessing the efficiency of the indicated siRNAs by measuring the expression levels of respective endogenous target in HEK293 cells. Data are mean + s.d. of n = 3 biologically independent samples.
**b)** Quantitative real-time PCR (qRT-PCR) assessing the expression levels of *AMOT130*, *AMOT-L1* and AMOT-L2 in HEK293 (left panel) and MCF10A (middle and right panels) treated with the indicated siRNA mixes directed against the three AMOT family members. Data are mean + s.d. of n = 3 biologically independent samples.
**c)** Representative γTubulin immunoblot of HEK293 cells treated with the indicated siRNAs. GAPDH serves as loading control.

**Supplementary Figure 3: Uncropped immunoblot images.**

Uncropped images of immunoblots, corresponding to those shown in the indicated figures. The immunoblot sections presented in the figures are indicated by dashed boxes.

---

**Extended Data Fig. 1:**
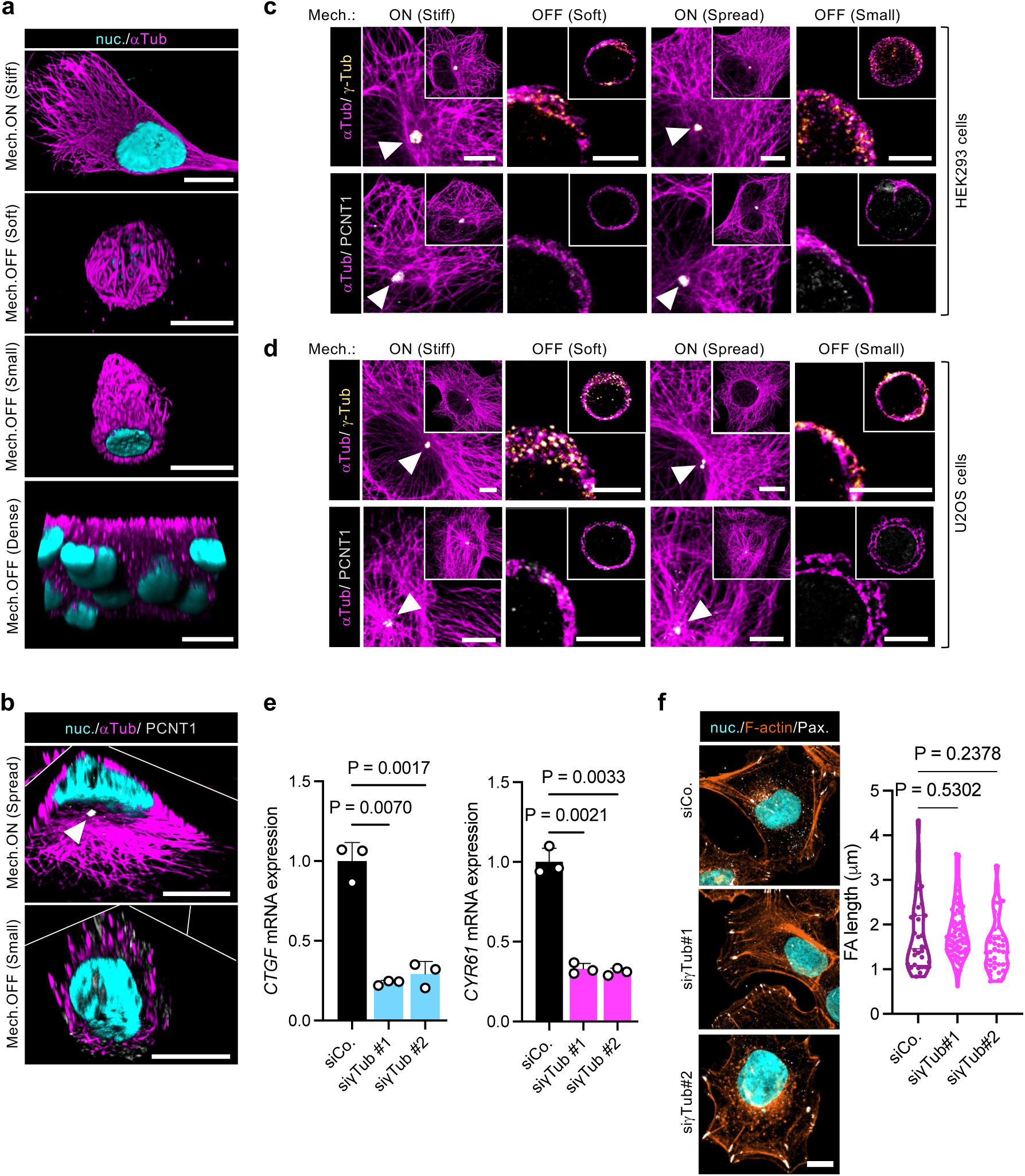
Microtubules (MTs) centrosomal organization serves as determinant of YAP/TAZ mechanotransduction, independently of F-actin and focal adhesions architecture. **a)** Representative 3D immunofluorescence reconstructions of MCF10A cells seeded the indicated Mech.ON versus Mech.OFF conditions. MTs were labelled by α-Tubulin staining (αTub, magenta) and nuclei were counterstained with Hoechst (cyan). Scalebar, 10μm. See also Supplementary Video 1 for an all-around view of the same cells. **b)** Representative immunofluorescence images (3D sections) of MCF10A cells seeded on Spread (unconfined, Mech.ON) versus Small (100 μm^2^ micropatterns, Mech.OFF) substrates. MTs were labelled by α-Tubulin staining (αTub, magenta), the centrosome was labelled by pericentrin (PCNT1, grey) and highlighted by white arrowhead. Nuclei were counterstained with Hoechst (cyan). Scale bar, 10μm. **c-d)** Representative immunofluorescence images of HEK293 cells (c) and U2OS cells (d) seeded in the indicated mechanical conditions. Images are magnifications of the insets shown in the upper right corner of each picture. MTs were labelled by α-Tubulin staining (αTub, magenta), γ-TURCs were labelled by γ-Tubulin staining (γTub, yellow), the centrosome was labelled by pericentrin (PCNT1, grey). γ-TURCs and centrosomes are highlighted by white arrowheads. Scale bar, 5μm. **e)** Quantitative real-time PCR (qRT-PCR) assessing the expression levels of the YAP/TAZ endogenous targets *CTGF* (left) and *Cyr61* (right) in HEK293 cells treated with the indicated siRNAs and seeded in mechano-ON conditions. Data are mean + s.d. of n = 3 biologically independent samples. **f)** Left: representative immunofluorescence images of HEK293 cells treated with the indicated siRNAs. F-actin was labelled by Phalloidin staining (F-actin, orange), focal adhesions were labelled by Paxillin staining (Pax, grey) and nuclei were counterstained with Hoechst (cyan). Scale bar, 10μm. Right: quantifications (n≥30) of focal adhesion length in cells treated as in left panels. *P* values were determined by one-way ANOVA with Welch’s correction (**e,f**).

**Extended Data Fig. 2:**
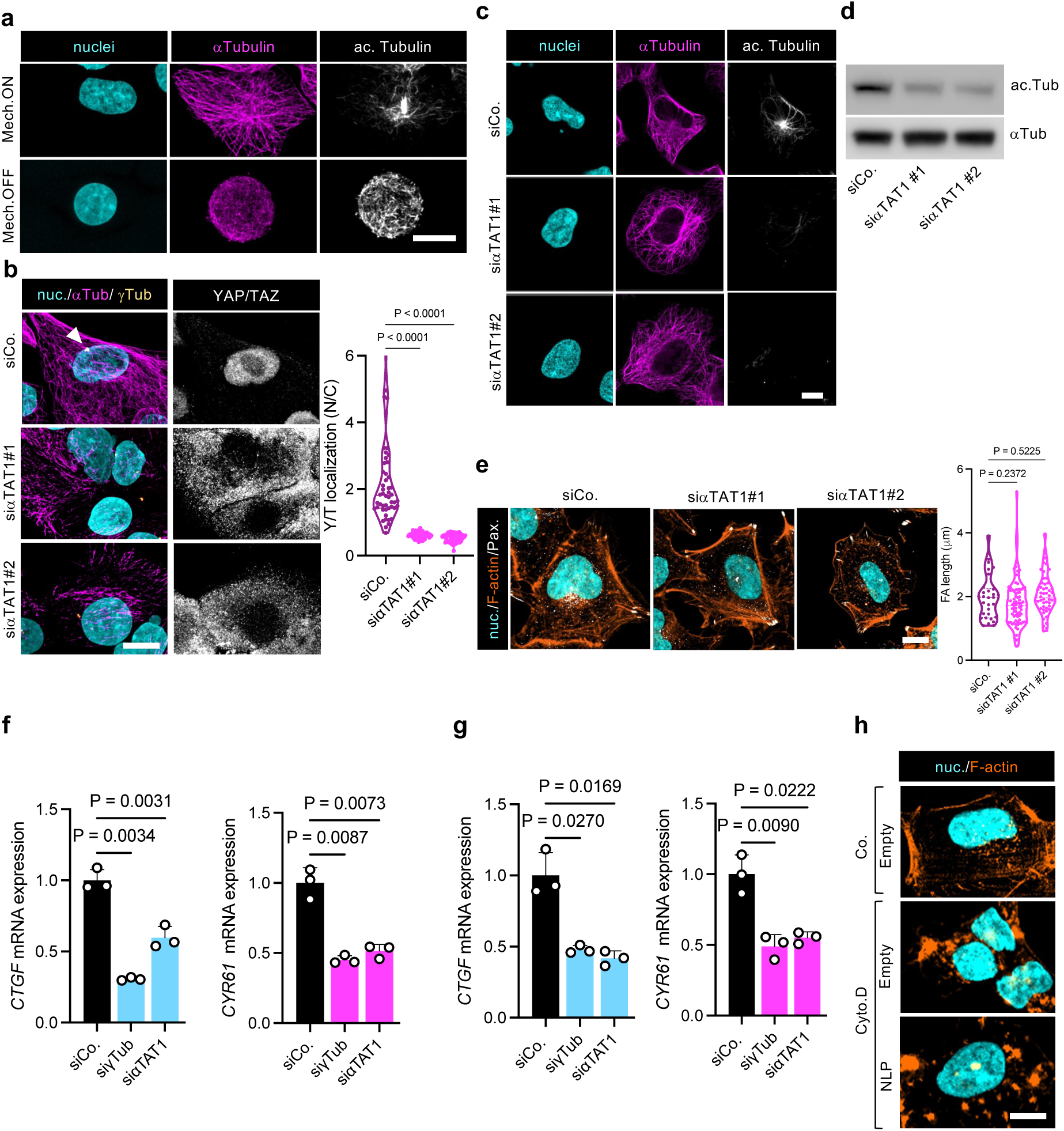
Microtubules (MTs) centrosomal organization is causal for YAP/TAZ regulation. **a)** Representative immunofluorescence images of MTs (αTub) and acetylated Tubulin (ac. Tubulin) in HEK293 cells seeded in the indicated mechanical conditions. Nuclei were counterstained with Hoechst (cyan). **b)** Left: Representative immunofluorescence images of MTs (αTub), γ-TURCs (γTub) and YAP/TAZ in HEK293 cells transfected with the indicated siRNAs. Nuclei were counterstained with Hoechst (cyan). White arrowhead indicates MTs convergence in the centrosomal MTOC in control cells. Scale bar, 10μm. Right: quantifications (n≥50) of YAP/TAZ nuclear-to-cytoplasmic subcellular localization (N/C) in cells treated as in left panels. **c)** Representative immunofluorescence images of MTs (αTub) and acetylated Tubulin (ac. Tubulin) in HEK293 cells seeded in Mech.ON conditions and treated with the indicated siRNAs. Nuclei were counterstained with Hoechst (cyan). **d)** Representative immunoblots of acetylated Tubulin (ac. Tub) in HEK293 cells seeded in mechano-ON conditions and treated with the indicated siRNAs. αTubulin (αTub) serves as loading control. **e)** Left: representative immunofluorescence images of HEK293 cells treated with the indicated siRNAs. F-actin was labelled by Phalloidin staining (F-actin, orange), focal adhesions were labelled by Paxillin staining (Pax, grey) and nuclei were counterstained with Hoechst (cyan). Scale bar, 10μm. Right: quantifications (n≥30) of focal adhesion length in cells treated as in left panels. **f,g)** Quantitative real-time PCR (qRT-PCR) assessing the expression levels of the YAP/TAZ endogenous targets *CTGF* (left) and *Cyr61* (right) in MCF10A (f) and U2OS (g) cells treated with the indicated siRNAs and seeded in mechano-ON conditions. Data are mean + s.d. of n = 3 biologically independent samples. **h)** Representative immunofluorescence of F-actin (orange, Phalloidin labelling) in HEK293 cells transduced with empty or NLP1-encoding lentiviruses and treated with Cytochalasin D (Cyto.D, 1μM for 2 hrs) or DMSO as negative control, as in Fig. 1g,h. Nuclei were counterstained with Hoechst (cyan). Scale bar, 5μm. *P* values were determined by one-way ANOVA with Welch’s correction (**e-g**).

**Extended Data Fig. 3:**
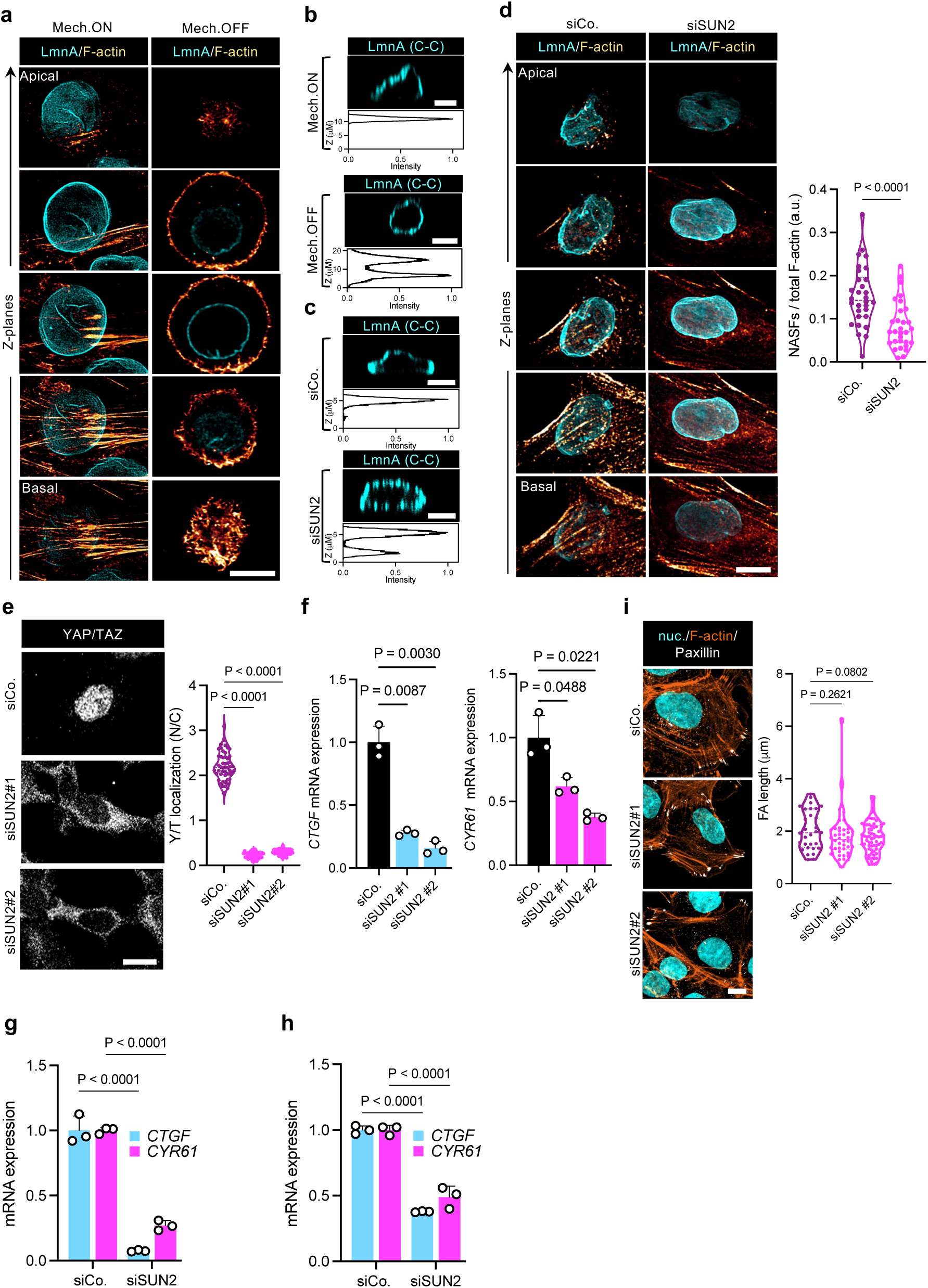
Loss of NE-NASFs tethering impairs YAP/TAZ mechano-activation. **a)** Representative Z-planes of immunofluorescence stacks of MCF10A cells seeded in mechano-ON (40 kPa) versus -OFF (0,7 kPa) conditions, showing NASFs contacting the basal side of the nuclear envelope of mechano-activated, but not of mechano-inhibited cells. The nuclear lamina was labelled by LaminA staining (cyan) and F-actin was labelled by Phalloidin staining (orange). Basal to apical planes are shown in bottom to top panels. Scale bar, 10μm. See also Fig. 1i for 3D reconstructions of cells seeded in the same conditions. **b)** Representative immunofluorescence YZ-sections of MCF10A cells seeded on stiff (Mech.ON) versus soft (Mech.OFF) hydrogels. Lamin was labelled by a conformation-sensitive LmnA antibody (LmnA C-C, cyan), which is unable to bind the basal side of the NE of mechano-ON cells due to LaminA stretching. The corresponding intensity profiles of LmnA conformation-sensitive staining along the Z-axis is shown below each immunofluorescence picture. Scale bar, 10μm. **c)** Top: representative immunofluorescence YZ-sections of MCF10A cells treated with the indicated siRNAs. A conformation-sensitive LmnA antibody (LmnA C-C, cyan), which is unable to bind stretched LaminA was used to show loss of basal nuclear tension in SUN2-depleted cells. The corresponding intensity profiles of LmnA conformation-sensitive staining along the Z-axis is shown below each immunofluorescence picture. Scale bar, 10μm. **d)** Left: Representative Z-planes of immunofluorescence stacks of MCF10A cells treated with indicated siRNAs, showing abolished NASFs formation in SUN2-depleted cells. The nuclear lamina was labelled by LaminA staining (cyan) and F-actin was labelled by Phalloidin staining (orange). Basal to apical planes are shown in bottom to top panels. Scale bar, 10μm. See also Fig. 1j for 3D sections of cells treated as in (**d**). Right: quantifications of NAFs/total F-actin ratios in the same cells (n≥30). **e)** Left: Representative immunofluorescence images of YAP/TAZ (grey) in HEK293 cells transfected with the indicated siRNAs. Scale bar, 10μm. Right: quantifications (n≥50) of YAP/TAZ nuclear-to-cytoplasmic subcellular localization (N/C) in cells treated as in left panels. **f)** Quantitative real-time PCR (qRT-PCR) assessing the expression levels of the YAP/TAZ endogenous targets *CTGF* (left) and *Cyr61* (right) in HEK293 cells treated with the indicated siRNAs and seeded in mechano-ON conditions. Data are mean + s.d. of n = 3 biologically independent samples. **g,h)** Quantitative real-time PCR (qRT-PCR) assessing the expression levels of the YAP/TAZ endogenous targets *CTGF* (left) and *Cyr61* (right) in MCF10A (g) and U2OS (h) cells treated with the indicated siRNAs and seeded in mechano-ON conditions. Data are mean + s.d. of n = 3 biologically independent samples. **i)** Loss of NE-NASFs tethering impairs YAP/TAZ mechano-activation, independently of F-actin and focal adhesions architecture. Left: representative immunofluorescence images of HEK293 cells treated with the indicated siRNAs. F-actin was labelled by Phalloidin staining (F-actin, orange), focal adhesions were labelled by Paxillin staining (Pax, grey) and nuclei were counterstained with Hoechst (cyan). Scale bar, 10μm. Right: quantifications (n≥30) of focal adhesion length in cells treated as in left panels. *P* values were determined by unpaired Student’s *t*-test with Welch’s correction (**d**), one-way ANOVA with Welch’s correction **(f,i)** or with two-way ANOVA (**g,h**) .

**Extended Data Fig. 4:**
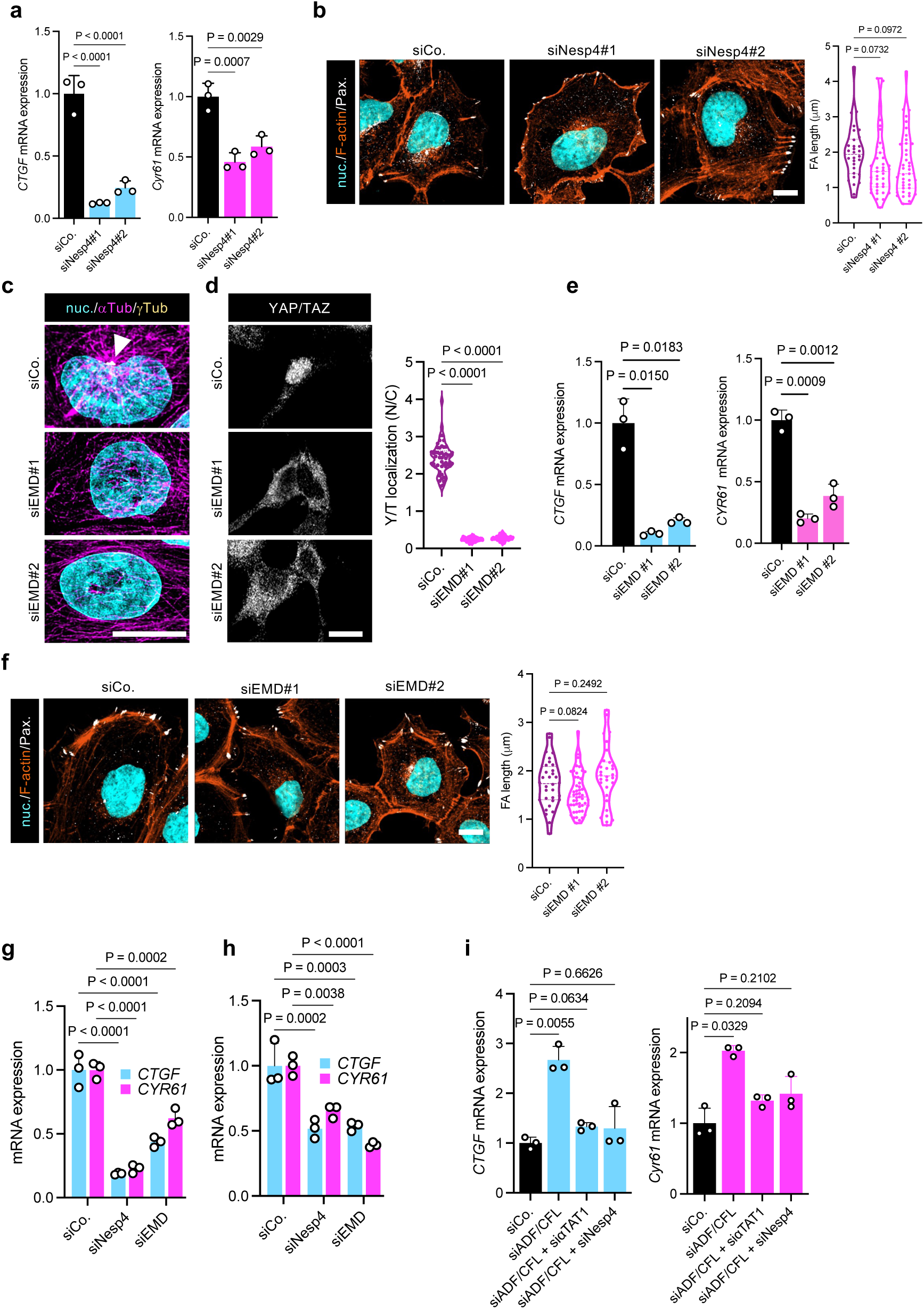
Loss of NE-MTs tethering impairs YAP/TAZ mechano-activation. **a)** Quantitative real-time PCR (qRT-PCR) assessing the expression levels of the YAP/TAZ endogenous targets *CTGF* (left) and *Cyr61* (right) in HEK293 cells treated with the indicated siRNAs and seeded in mechano-ON conditions. Data are mean + s.d. of n = 3 biologically independent samples. **b)** Left: representative immunofluorescence images of HEK293 cells treated with the indicated siRNAs. F-actin was labelled by Phalloidin staining (F-actin, orange), focal adhesions were labelled by Paxillin staining (Pax, grey) and nuclei were counterstained with Hoechst (cyan). Scale bar, 10μm. Right: quantifications (n≥30) of focal adhesion length in cells treated as in left panels. **c)** Representative immunofluorescence images of MTs (αTub) and γ-TURCs (γTub) in HEK293 cells transfected with the indicated siRNAs. White arrowhead highlights MTs convergence in a perinuclear MTOC only in control (siCo.) cells. Nuclei were counterstained with Hoechst (cyan). Scale bar, 10μm. **d)** Left: representative immunofluorescence images of YAP/TAZ (grey) in HEK293 cells transfected with the indicated siRNAs. Scale bar, 10μm. Right: quantifications (n≥50) of YAP/TAZ nuclear-to-cytoplasmic subcellular localization (N/C) in cells treated as in left panels. **e)** Quantitative real-time PCR (qRT-PCR) assessing the expression levels of the YAP/TAZ endogenous targets *CTGF* (left) and *Cyr61* (right) in HEK293 cells treated with the indicated siRNAs and seeded in mechano-ON conditions. Data are mean + s.d. of n = 3 biologically independent samples. **f)** Loss of NE-MTs tethering impairs YAP/TAZ mechano-activation, independently of F-actin and focal adhesions architecture. Left: representative immunofluorescence images of HEK293 cells treated with the indicated siRNAs. F-actin was labelled by Phalloidin staining (F-actin, orange), focal adhesions were labelled by Paxillin staining (Pax, grey) and nuclei were counterstained with Hoechst (cyan). Scale bar, 10μm. Right: quantifications (n≥30) of focal adhesion length in cells treated as in left panels. **g,h)** Quantitative real-time PCR (qRT-PCR) assessing the expression levels of the YAP/TAZ endogenous targets *CTGF* (left) and *Cyr61* (right) in MCF10A (g) and U2OS (h) cells treated with the indicated siRNAs and seeded in mechano-ON conditions. Data are mean + s.d. of n = 3 biologically independent samples. **i)** Quantitative real-time PCR (qRT-PCR) assessing the expression levels of the YAP/TAZ endogenous target *CTGF* (left) and *Cyr61* (right) in HEK293 cells treated with the indicated siRNAs and seeded in mechano-OFF conditions. Data are mean + s.d. of n = 3 biologically independent samples. *P* values were determined by one-way ANOVA with Welch’s correction **(a,b,d-g,i)**, or with two-way ANOVA **(g,h)**.

**Extended Data Fig. 5:**
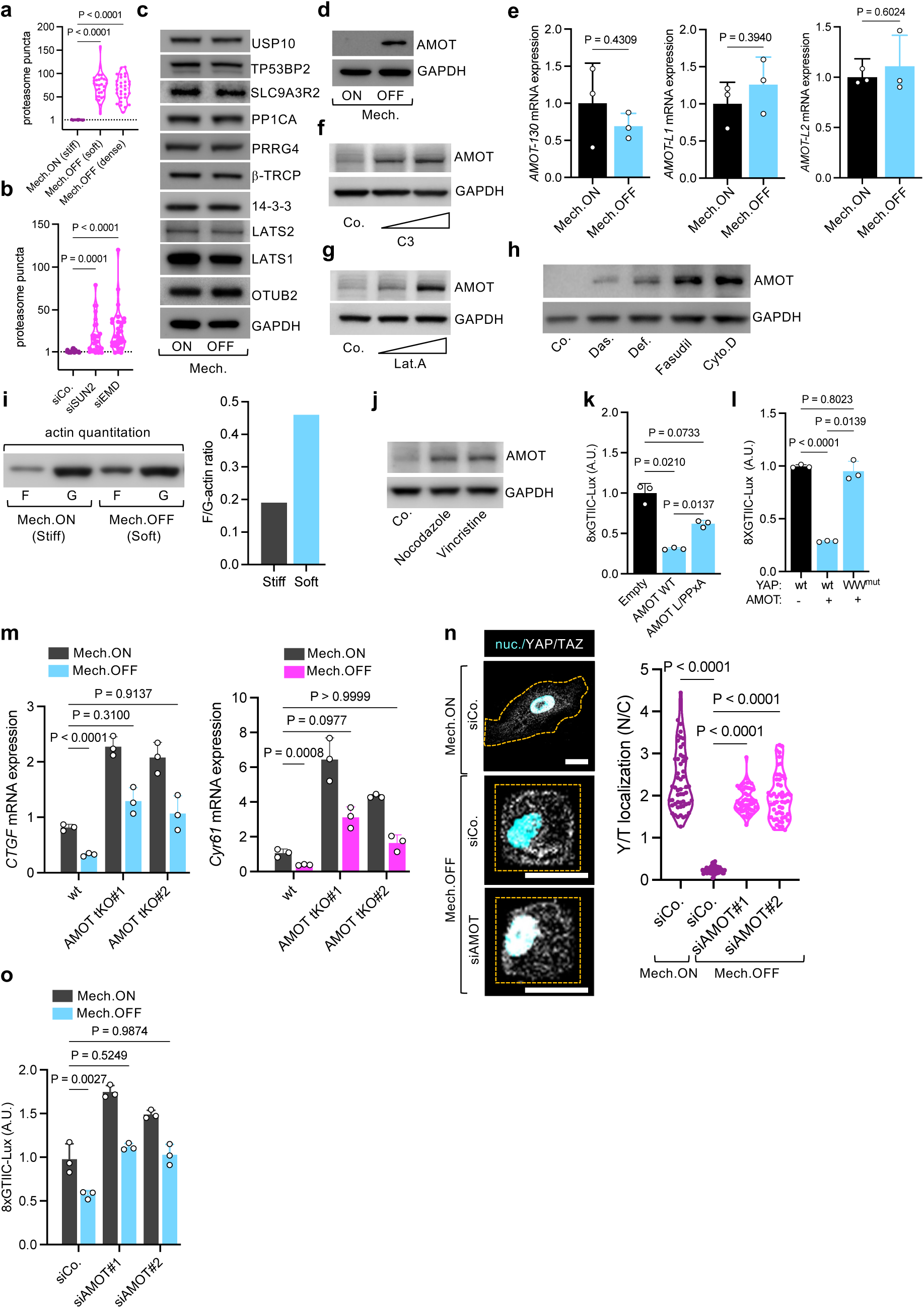
AMOT acts as cytoplasmic mechanical rheostat to control YAP/TAZ activity. **a)** Quantifications (n≥30) of proteasome (20-S staining) in cells seeded as in Fig. 2a. Preferential accumulation of the proteasome in a single perinuclear area is set to 1 in mechano-ON conditions. Conversely, cells seeded in mechano-OFF displayed multiple proteasome signals scattered throughout the cytoplasm. **b)** Quantifications (n≥30) of proteasome (20-S staining) in cells seeded as in Fig. 2b. Preferential accumulation of the proteasome in a single perinuclear area is set to 1 in siCo. conditions. Conversely, cells depleted of SUN2 or Emerin displayed multiple proteasome signals scattered throughout the cytoplasm. **c)** Representative immunoblot of YAP/TAZ cytoplasmic interactors in HEK293 cells seeded in mechano-ON (stiff) versus mechano-OFF (0.7 kPa hydrogels) conditions. GAPDH serves as loading control. See also Supplementary Table 1 for a summary of these results. **d)** Representative AMOT immunoblot of HEK293 cells seeded in mechano-ON (sparse) versus mechano-OFF (dense) conditions. GAPDH serves as loading control. **e)** Quantitative real-time PCR (qRT-PCR) showing that the endogenous expression levels of AMOT-130 and AMOT-L1 (HEK293 cells) and AMOT-L2 (MCF10A cells) are unaffected by Mech.ON (stiff) versus Mech.OFF (0.7 kPa hydrogels) conditions. Data are mean + s.d. of n = 3 biologically independent samples. **f)** Representative AMOT immunoblot of HEK293 cells treated with increasing doses of the Rho inhibitor 1 (Exoenzyme C3 Transferase, 5 μg ml^-1^ and 10 μg ml^-1^, 4 hours). GAPDH serves as loading control. **g)** Representative AMOT immunoblot of HEK293 cells treated with increasing doses of LatrunculinA (Lat.A, 343 nM and 686 nM, 2 hours). GAPDH serves as loading control. **h)** Representative AMOT immunoblot of HEK293 cells treated with the indicated mechano-inhibitory drugs (Das., Dasatinib 100nM; Def., Defactinib 5 μM; Fasudil 10 μM; Cyto.D, Cytochalasin D 1μM for 2 hours). GAPDH serves as loading control. **i)** Left: representative immunoblot showing quantitation of filamentous (F) versus free globular (G) actin in MCF10A cells seeded on stiff (40kPa) versus soft (0.7 kPa) hydrogels. Right: quantitation of F/G-actin ratios of the samples shown on the left (see methods). **j)** Representative AMOT immunoblot of HEK293 cells treated with the indicated inhibitors of microtubule polymerization (Nocodazole 5μM; Vincristine 5μM, for 1 hour). GAPDH serves as loading control. **k)** Luciferase assay of HEK293 cells seeded in mechano-ON conditions and transfected with a synthetic reporter for YAP/TAZ/TEAD-dependent transcription (8xGTIIC-Lux) and with the indicated AMOT mutants. Data are mean + s.d. of n = 3 biologically independent samples. **l)** Luciferase assay of HEK293 cells treated with siRNAs targeting YAP/TAZ and reconstituted with either YAPwt of YAP-WW mutant constructs (see methods). Cells were concomitantly transiently transfected with AMOT-expressing or empty vectors, and with a synthetic reporter for YAP/TAZ/TEAD-dependent transcription (8xGTIIC-Lux), and after 24 hours seeded in mechano-ON conditions. Data are mean + s.d. of n = 3 biologically independent samples. **m)** Quantitative real-time PCR (qRT-PCR) assessing the expression levels of the YAP/TAZ endogenous targets *CTGF* (left) and *Cyr61* (right) in control (wt) or AMOT tKO M2 (see methods) cells seeded in Mech.ON (stiff) versus Mech.OFF (0.7 kPa hydrogels) conditions. Data are mean + s.d. of n = 3 biologically independent samples. **n)** Left: representative immunofluorescence images of YAP/TAZ in hMSCs treated with the indicated siRNAs and seeded in Mech.OFF (1000 μm^2^ micropatterns) versus Mech.ON (unconfined) conditions. Nuclei were counterstained with Hoechst (cyan). Scale bar, 20μm. Cell borders are outlined by orange dashes. Right: quantifications (n≥50) of YAP/TAZ nuclear-to-cytoplasmic subcellular localization (N/C) in cells seeded as in left panels. **o)** Luciferase assay of HEK293 cells treated with the indicated siRNAs, seeded in mechano-ON versus mechano-OFF conditions and transfected with a synthetic reporter for YAP/TAZ/TEAD-dependent transcription (8xGTIIC-Lux). Data are mean + s.d. of n = 3 biologically independent samples. *P* values were determined by unpaired Student’s *t*-test with Welch’s correction (**a**,**e**), one-way ANOVA with Welch’s correction **(b, k, l,n)**, or with two-way ANOVA (**m, o**).

**Extended Data Fig. 6:**
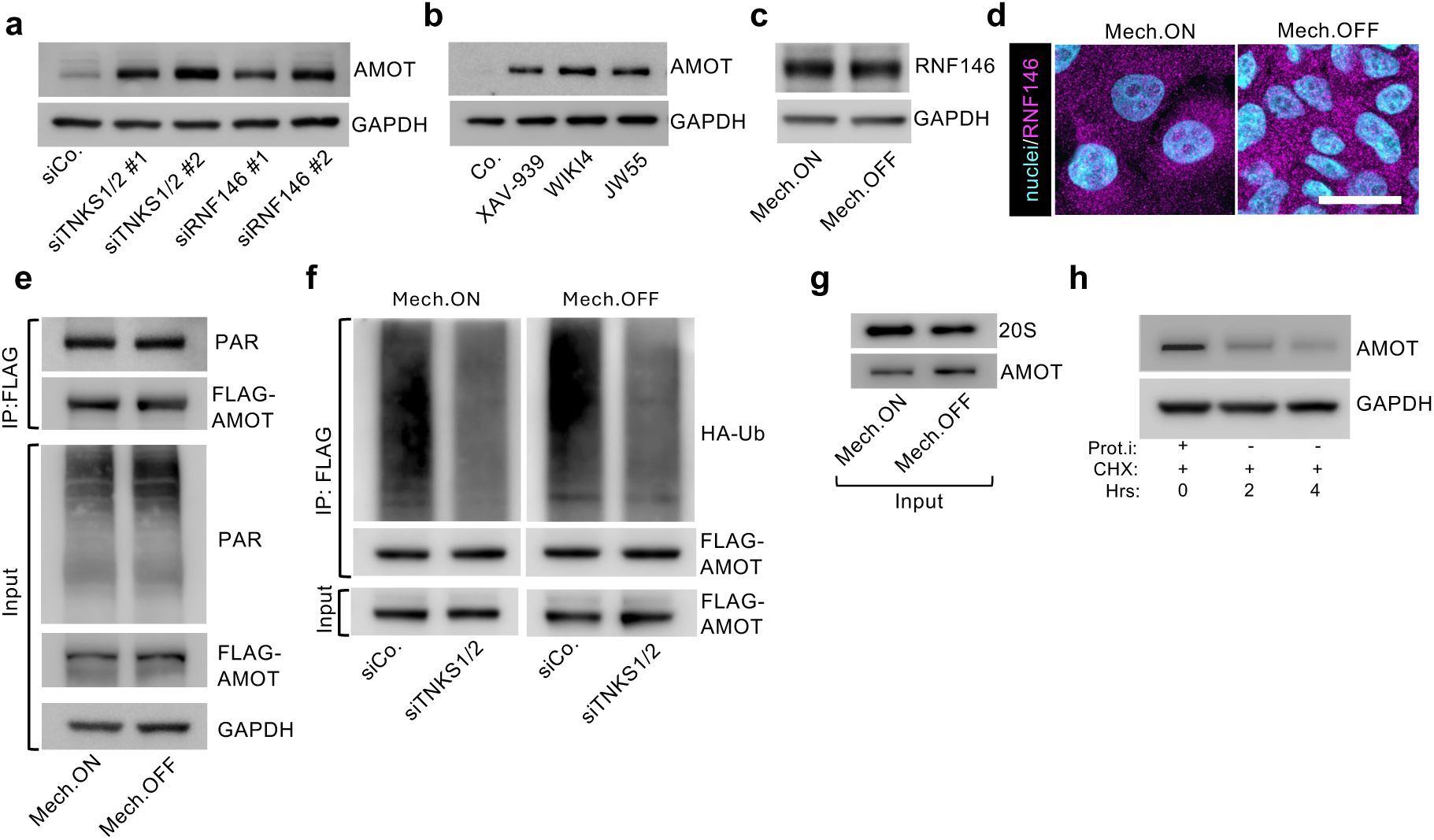
AMOT is constitutively tagged for proteasomal degradation under both mechano-ON and mechano-OFF conditions. **a)** Representative AMOT immunoblot of HEK293 cells treated with siRNAs targeting the poly-ADP-ribosyltransferase TNKS1/2 or the E3 ubiquitin ligase RNF146 and seeded in mechano-ON conditions. GAPDH serves as loading control. **b)** Representative AMOT immunoblot of HEK293 cells treated with the indicated independent Tankyrase1/2 inhibitors (10 μM, 24 hours). GAPDH serves as loading control. **c)** Representative RNF146 immunoblot of HEK293 cells seeded in Mech.ON (sparse) versus Mech.OFF (dense) conditions. GAPDH serves as loading control. **d)** Representative immunofluorescence images of RNF146 in HEK293 cells seeded in Mech.ON (spread) versus Mech.OFF conditions (dense). Nuclei were counterstained with Hoechst (cyan). Scale bar, 25 μm. **e)** Representative PARylation assay (see methods) in HEK293 cells seeded in Mech.ON (sparse) versus Mech.OFF (dense) conditions and treated with proteasome inhibitors showing that AMOT PARylation levels are unaffected by the mechanical state of the cell. GAPDH serves as loading control for inputs. **f)** Representative Ubiquitination assays (see methods) in HEK293 cells seeded in Mech.ON (sparse) versus Mech.OFF (dense) conditions and treated the indicated siRNAs, showing that Tankyrase1/2-deopendent AMOT poly-ubiquitination occurs irrespective of the the mechanical state of the cell. **g)** Inputs for the co-immunoprecipitation experiment shown in Fig. 3c. **h)** Cycloheximide (CHX) pulse-and-chase experiment (see methods) in HEK293 cells showing that in mechano-ON conditions, AMOT is rapidly degraded after proteasome inhibitor (Prot.i) washout. Cells were treated with CHX 50 μg/mL. GAPDH serves as loading control.

**Extended Data Fig. 7:**
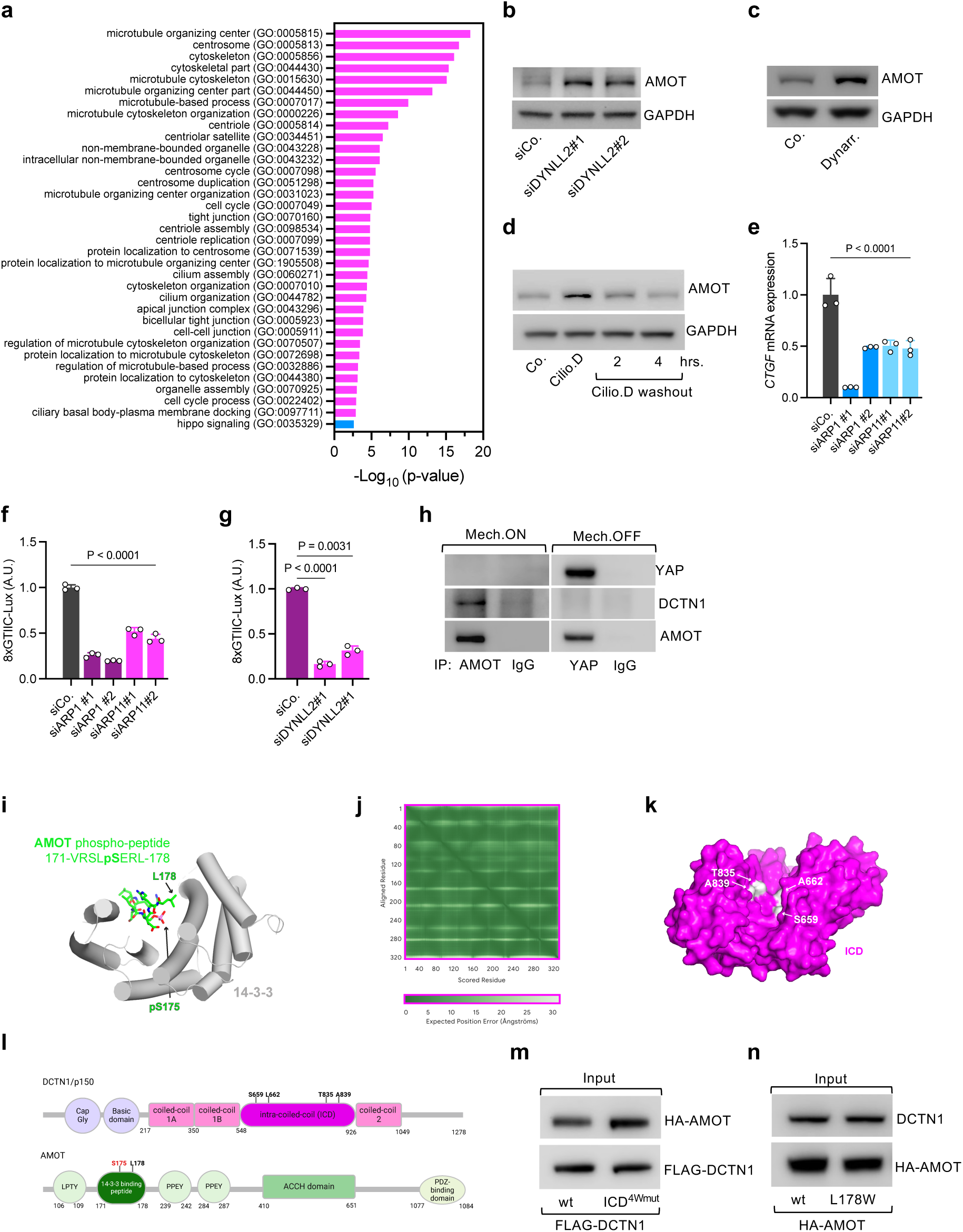
AMOT delivery to proteasomal condensates by dynein-mediated retrograde transport. **a)** Graph depicting the most significant GO terms emerging from the Gene Ontology analyses of AMOT proximal proteins, as reported in a publicly available dataset^38^. A full list of retrieved AMOT proximal proteins and associated significant GO terms (-Log_10_[p-value]) ≥ 2.6) is provided in Supplementary Table 2. **b)** Representative AMOT immunoblot of HEK293 cells seeded in mechano-ON conditions and treated with the indicated siRNAs. GAPDH serves as loading control. **c)** Representative AMOT immunoblot of HEK293 cells seeded in mechano-ON conditions and treated with DMSO (Co.) or Dynarrestin (10 μM, 6 hours). GAPDH serves as loading control. **d)** Representative AMOT immunoblot of HEK293 cells seeded in mechano-ON conditions and treated with DMSO (Co.) or Ciliobrevin D (Cilio.D 10 μM, 5 hours). Cilio.D treatment was followed by 2 or 4 hours of washout. GAPDH serves as loading control. **e)** Quantitative real-time PCR (qRT-PCR) assessing the expression levels of the YAP/TAZ endogenous target *CTGF* in HEK293 cells treated with the indicated siRNAs and seeded in mechano-ON conditions. Data are mean + s.d. of n = 3 biologically independent samples. **f-g)** Luciferase assay of HEK293 cells treated with the indicated siRNAs, seeded in mechano-ON conditions and transfected with a synthetic reporter for YAP/TAZ/TEAD-dependent transcription (8xGTIIC-Lux). Data are mean + s.d. of n = 3 biologically independent samples. **h)** Pulldown of endogenous AMOT (left) or endogenous YAP (right) from proteasome-inhibited HEK293T cell seeded in Mech.ON versus Mech.OFF conditions, showing that AMOT bound to DCTN1 in Mech.ON cells is unable to bind to YAP. IgG pulldown serves as negative control. **i)** Cartoon representations of the X-ray structure (PDB code 7nma) ^43^ of the 14-3-3 domain (grey) in complex with a short AMOT peptide phosphorylated at S175 (pS175). The AMOT peptide is shown as stick representation in green with pS175 and L178 highlighted. **j)** AlphaFold2 predicted alignment error (PAE) plot ^84^ for the complex shown in Fig. 3g, indicating that the ICD structure is predicted with very high confidence. **k)** DCTN1 ICD as surface representation with highlighted in white the four residues (S659, A662, T835, A839) lining the putative binding cleft. These have been replaced by tryptophans for co-immunoprecipitation (co-IP) assays of Fig. 3h (see main text for details). **l)** Schematic illustrations of the domain structures and motifs of DCTN1/p150 (top) and AMOTp130 (bottom). **m)** Representative immunoblots of the inputs of the pulldown experiment shown in Fig. 3h. **n)** Representative immunoblots of the inputs of the pulldown experiment shown in Fig. 3i. *P* values were determined by one-way ANOVA with Welch’s correction (**e-g**).

**Extended Data Fig. 8:**
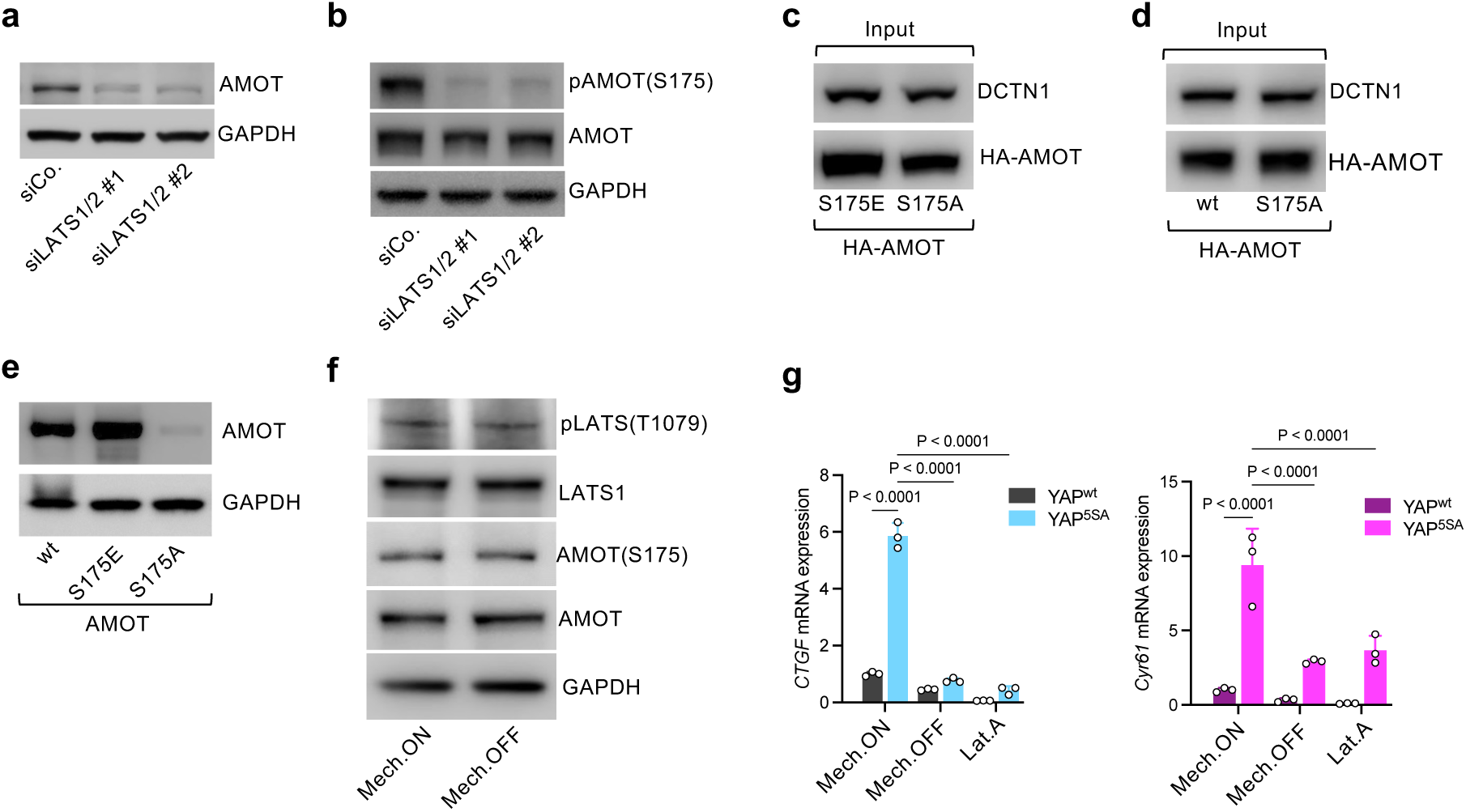
Hippo/LATS signaling indirectly feeds on YAP/TAZ mechanoregulation through AMOT. **a)** Representative AMOT immunoblot of HEK293 cells treated with the indicated siRNAs and seeded in mechano-OFF conditions. GAPDH serves as loading control. **b)** Representative phospho-AMOT-S175 immunoblot of HEK293 cells transfected with the indicated siRNAs, seeded in mechano-ON conditions and treated with proteasome inhibitor (Lactacystin, 10 μM, 8 hours). GAPDH serves as loading control. **c)** Representative immunoblots of the inputs of the pulldown experiment shown in Fig. 4b. **d)** Representative immunoblots of the inputs of the pulldown experiment shown in Fig. 4c. **e)** Representative AMOT immunoblots of HEK293 cells transiently transfected with the indicated AMOT mutants. GAPDH serves as loading control. **f)** Representative immunoblots of HEK293 cells seeded in Mech.ON (stiff) versus Mech.OFF (0.7 kPa hydrogels) conditions treated with proteasome inhibitor (Lactacystin, 10 μM, 8 hours), showing that AMOT-S175 and LATS1-T1079 phosphorylation is unaffected by cell mechanics. GAPDH serves as loading control. **g)** Quantitative real-time PCR (qRT-PCR) assessing the expression levels of the YAP/TAZ endogenous targets *CTGF* (left) and *Cyr61* (right) in YAPwt or YAP5SA reconstituted MCF10A cells (see methods) seeded in Mech.ON (sparse), Mech.OFF (dense) conditions or treated with Lat.A (343 nM, 15 hours). Data are mean + s.d. of n = 3 biologically independent samples. *P* values were determined by two-way ANOVA (**g**).

**Extended Data Fig. 9:**
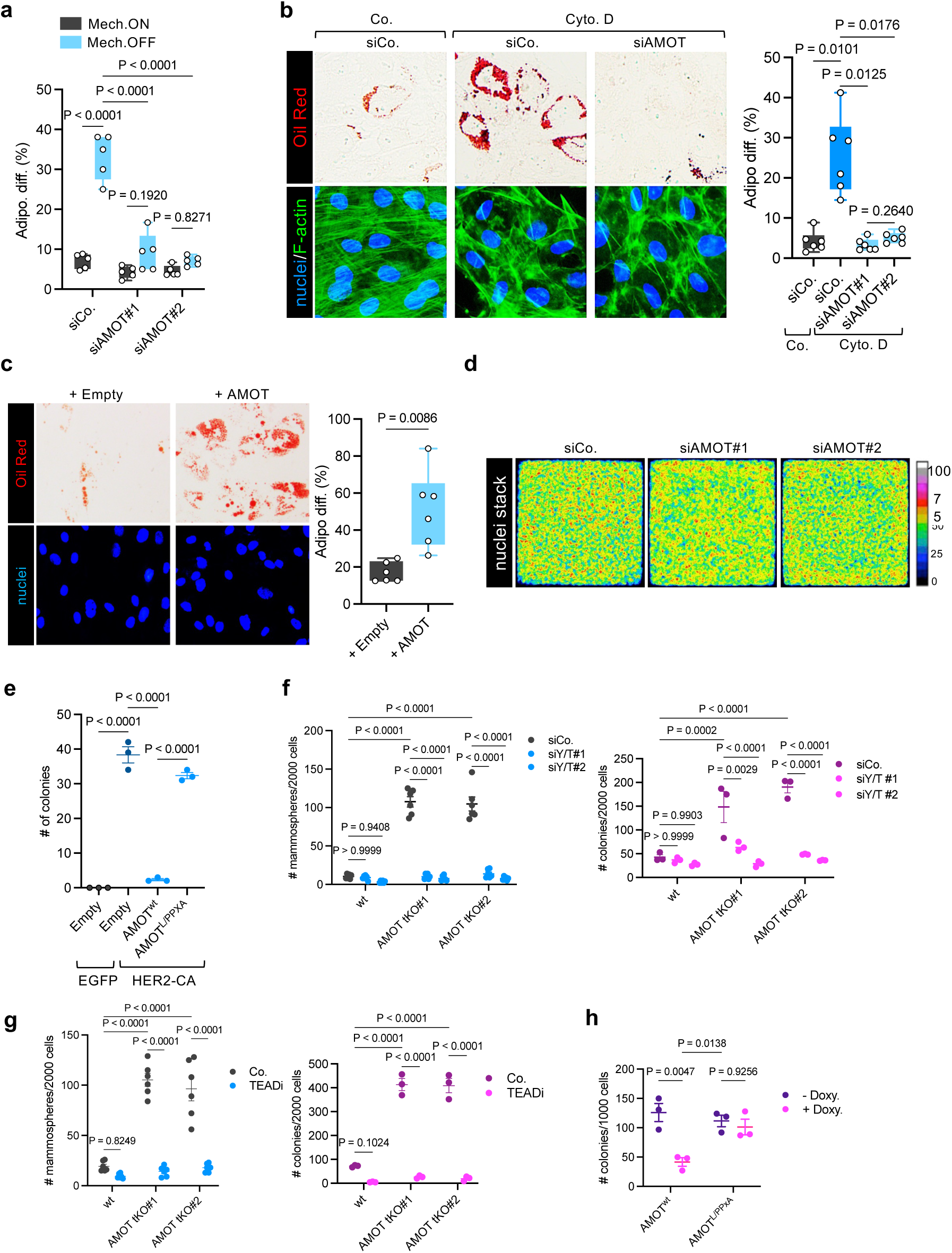
AMOT acts as mechanical rheostat to control biological responses driven by YAP/TAZ. **a)** Quantifications of the percentage of adipogenic differentiation (based on Oil Red staining) of hMSCs seeded on spread (unconfined, Mech.ON) or small (1000 μm^2^, Mech.OFF) micropatterns and treated with the indicated siRNAs, as in Fig. 5a. Data are representative of n= 5 biologically independent experiments, with n ≥ 100 cells quantified each. **b)** Left: Representative Oil Red staining of hMSCs transfected with the indicated siRNAs, seeded in mechano-ON conditions and treated with DMSO (Co.) or with Cytochalasin D (Cyto.D, see methods). To verify Cyto.D effectiveness, F-actin was labelled with phalloidin (F-actin, green). Nuclei were counterstained with Hoechst (blue). Right: quantifications of the percentage of adipogenic differentiation (based on Oil Red staining) of hMSCs treated as in left panels. Data are representative of n= 6 biologically independent experiments, with n ≥ 100 cells quantified each. **c)** Left: Representative Oil Red staining of hMSCs seeded in mechano-ON conditions, transduced with Empty or AMOT-encoding lentiviral vectors and subjected to adipogenic differentiation protocol (see methods). Nuclei were counterstained with Hoechst (blue). Right: quantifications of the percentage of adipogenic differentiation (based on Oil Red staining) of hMSCs treated as in left panels. Data are representative of n= 6 biologically independent experiments, with n ≥ 100 cells quantified each. **d)** Representative colorimetric stacked images of Hoechst counterstained nuclei of MCF10A cells treated with the indicated siRNAs and seeded as cell monolayers of defined dimensions as in Fig. 5b. The color scale indicates the density of Hoechst labelled nuclei, showing a uniform distribution of cells within the monolayers in all the experimental conditions. **e)** Quantifications of colonies formed by human mammary luminal differentiated cells undergoing oncogenic reprograming when transduced with lentiviral vectors encoding for the indicated factors, as in Fig. 5d: HER-CA, constitutive active HER2 mutant (see methods); AMOT^L/PPXA^, AMOT point mutant in the three L/PPXY YAP-binding motifs (see methods). **f)** Left: Quantifications of mammospheres formed by control (wt) or AMOT tKO MII cells treated with the indicated siRNAs. Data are representative of n = 6 biological independent samples. Right: Quantifications of soft agar colonies formed by control (wt) or AMOT tKO MII cells treated with the indicated siRNAs. Data are representative of n = 3 biological independent samples **g)** Left: Quantifications of mammospheres formed by control (wt) or AMOT tKO MII cells treated with the TEAD inhibitor VT107 (see methods), or with DMSO (Co.). Data are representative of n = 6 biological independent samples. Right: Quantifications of soft agar colonies formed by control (wt) or AMOT tKO MII cells treated with the with the TEAD inhibitor VT107 (see methods), or with DMSO (Co.). Data are representative of n = 3 biological independent samples **h)** Quantifications of soft agar colonies formed by MDA-MB-231 cells transduced with lentiviral vectors encoding for the indicated doxycycline (Doxy)-inducible AMOT mutant constructs. Data are representative of n = 3 biological independent samples. *P* values were determined by two-way ANOVA (**a,f-h**), or by one-way ANOVA with Welch’s correction (**e**), or by unpaired Student’s *t*-test with Welch’s correction (**c**).

**Extended Data Fig. 10:**
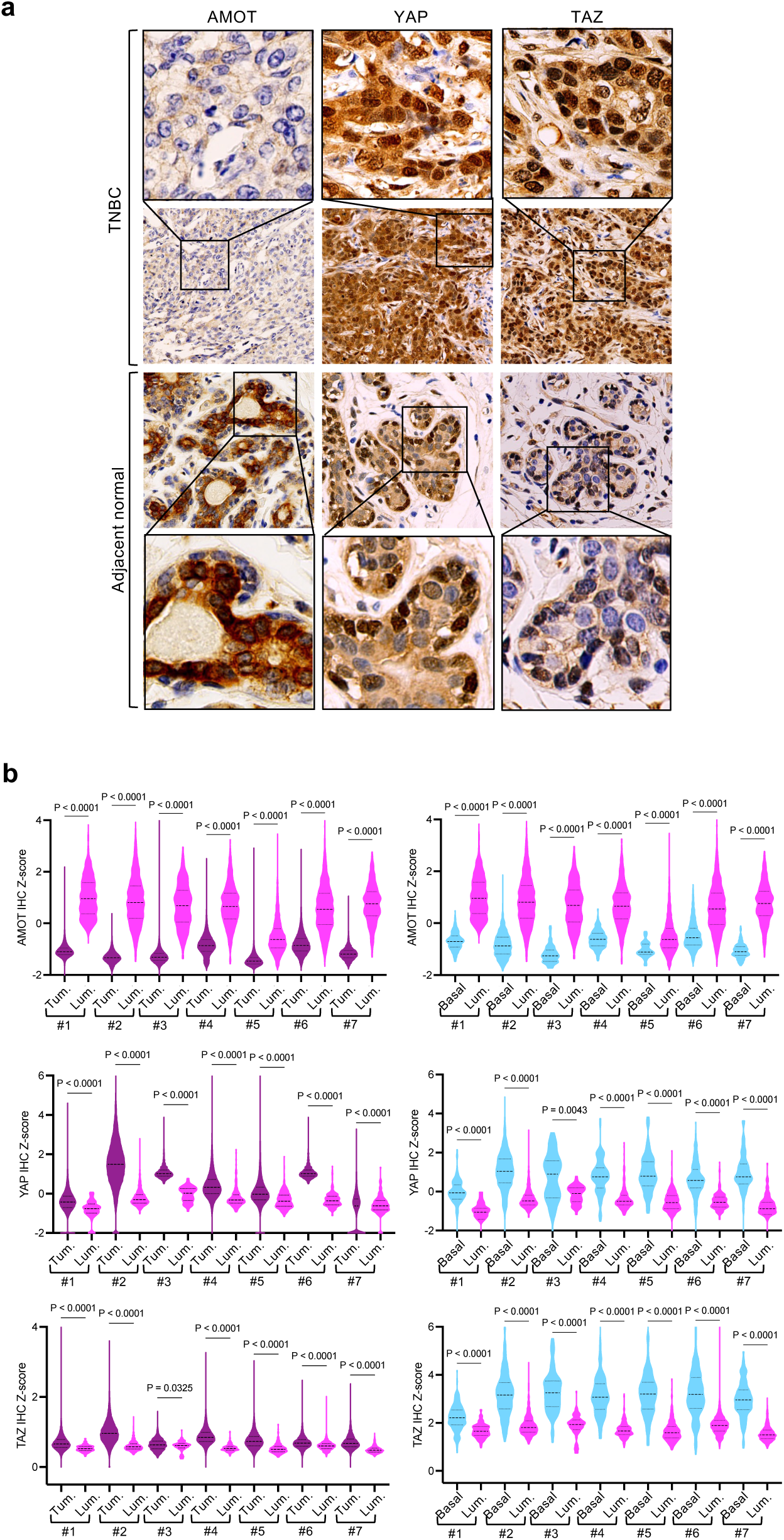
Role of AMOT as candidate tumor suppressor. **a)** Representative large field immunohistochemical pictures of AMOT, YAP and TAZ proteins in chemo-naïve human triple negative breast cancer (TNBC) samples or adjacent normal mammary tissue. The enlargements (top and bottom panels) are the same pictures shown in Fig. 5g. Data are representative of n=7 independent patient samples. **b)** Digital-pathology based quantifications (Z-scores, see methods) of AMOT abundance and YAP/TAZ nuclear-to-cytoplasmic subcellular localization based on immunohistochemical signals in the indicated cell populations (Tumor, Tum.; adjacent normal luminal, Lum.; and adjacent normal basal mammary cells, Basal) in independent TNBC samples (#1-#7), as shown in representative pictures of Fig. 5g and Extended Data Fig. 10a. n ≥ 100000 cells were quantified for each sample. *P* values were determined by unpaired Student’s *t*-test with Welch’s correction (**b**).

